# Discovery and characterisation of an amidine-containing ribosomally-synthesised peptide that is widely distributed in nature

**DOI:** 10.1101/2020.05.04.076059

**Authors:** Alicia H. Russell, Natalia M. Vior, Edward S. Hems, Rodney Lacret, Andrew W. Truman

## Abstract

Ribosomally synthesised and post-translationally modified peptides (RiPPs) are a structurally diverse class of natural product with a range of bioactivities. Genome mining for RiPP biosynthetic gene clusters (BGCs) is often hampered by poor detection of the short precursor peptides that are ultimately modified into the final molecule. Here, we utilise a previously described genome mining tool, RiPPER, to identify novel RiPP precursor peptides near YcaO-domain proteins, enzymes that catalyse various RiPP post-translational modifications including heterocyclisation and thioamidation. Using this dataset, we identified a novel, diverse and highly conserved family of RiPP BGCs spanning over 230 species of Actinobacteria and Firmicutes. A representative BGC from *Streptomyces albus* J1074 was characterised, leading to the discovery of streptamidine, a novel-amidine containing RiPP. This highlights the breadth of unexplored natural products with structurally rare features, even in model organisms.

## INTRODUCTION

Microorganisms produce an array of natural products (NPs) with broad biological activities. The phylum Actinobacteria is a particularly prominent source of NPs that have been utilised as antimicrobial drugs. Since the genome of the model actinobacterium *Streptomyces coelicolor* A3(2) was sequenced in 2002,^[1]^ it has been widely shown that bacteria are capable of producing many more NPs than are currently known, due to the abundance of uncharacterised biosynthetic gene clusters (BGCs) present in microbial genomes.^[2, 3]^ Ribosomally synthesised and post-translationally modified peptides (RiPPs) are a large and growing class of structurally diverse NPs. RiPPs are produced from a ribosomally synthesised precursor peptide that is typically comprised of a leader region and a core region. Post-translational modifications are installed onto the core region of the precursor peptide by a series of RiPP tailoring enzymes (RTEs),^[4, 5]^ which introduce structural diversity and complexity. The leader peptide is usually proteolytically removed as a late-stage step in RiPP biosynthesis.^[4, 6]^

Whilst genome mining is a popular approach to identify uncharacterised BGCs, the identification of novel RiPP BGCs is particularly challenging. This is because the small precursor peptides that are ultimately transformed into the final product are often not annotated in genomes, and unlike with other natural product classes such as polyketides, terpenes and non-ribosomal peptides, the short biosynthetic pathways for RiPPs lack universally shared features.^[7]^ Whilst some specific genome mining tools for RiPPs have been developed,^[8–11]^ many of these tools rely on the identification of homology to known RiPP classes. Therefore, the opportunity to identify truly novel RiPP families, and subsequent untapped structural complexity, might be missed. In the last two decades, hundreds of thousands of bacterial genomes have been sequenced, but their biosynthetic capacities have not been fully explored. The use of more bespoke genome mining tools therefore represents an important opportunity to identify cryptic and uncharacterised BGCs.

One of the most widespread families of proteins associated with RiPP biosynthesis are YcaO-domain proteins, which are ATP-dependent enzymes found in both bacteria and archaea,^[12]^ and have been shown to catalyse various post-translational modifications of RiPPs (Figure 1). This includes the installation of oxazoline and thiazoline heterocycles onto the precursor peptide backbone, where cyclodehydration is catalysed by the YcaO-domain in cooperation with a protein homologous to an E1 ubiquitin-activation enzyme or an “Ocin-ThiF-like” protein.^[13]^ YcaO proteins have also been demonstrated to catalyse the formation of amidine rings in bottromycin^[14]^ and klebsazolicin,^[15]^ and can also function with a TfuA-domain protein to introduce thioamide bonds into RiPPs such as thiopeptin^[16]^ and the thiovarsolins.^[7, 17]^ Over 9,000 YcaO-domain proteins have been bioinformatically identified in GenBank, but the function of the majority of these remain unknown.^[18]^

**Figure 1.**
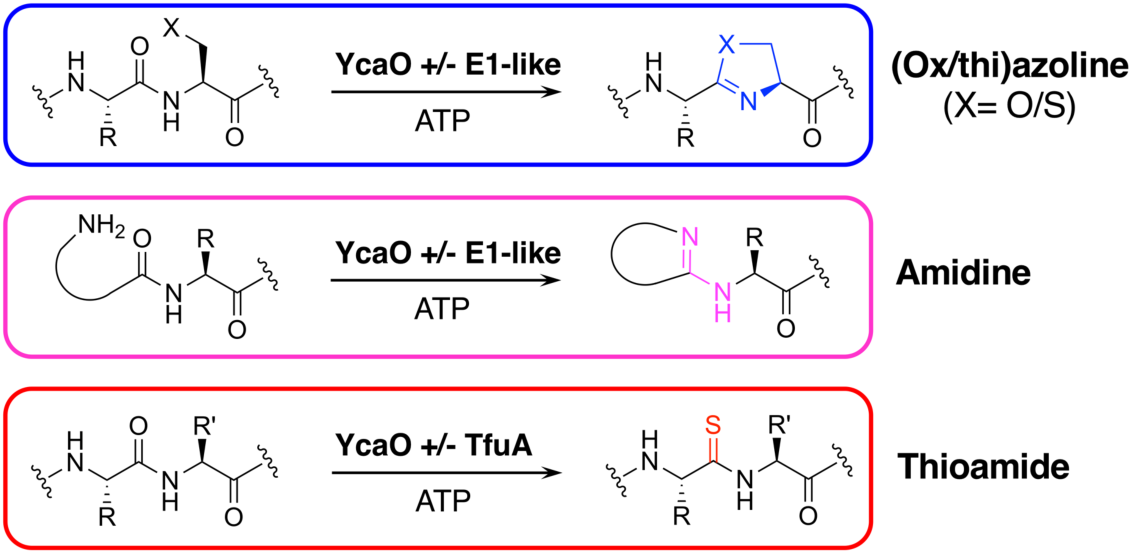
Reactions catalysed by YcaO-domain proteins.

We have previously reported a RiPP genome mining tool, RiPPER^[7]^ (RiPP Precursor Peptide Enhanced Recognition), which identifies precursor peptides without the need for information about RiPP structural class. RiPPER captures surrounding DNA regions of putative RTEs and searches for short open reading frames (ORFs) that might encode RiPP precursor peptides. In this study, we use RiPPER to identify precursor peptides encoded near all standalone YcaO-domain proteins in Actinobacteria. This identified a large family of novel and diverse RiPP BGCs that span over 230 bacterial species. The sequence variation of both the identified precursor peptides, as well as the associated RTEs, suggests that these BGCs produce structurally distinct molecules with variable post-translational modifications. We characterised an exemplar of this new RiPP BGC family from the model actinobacterium, *Streptomyces albus* J1074 (recently reclassified as *Streptomyces albidoflavus*),^[19]^ which led to the discovery of streptamidine, a novel and structurally rare amidine-containing molecule. The prevalence of this RiPP family highlights that we are still scratching the surface of the huge biosynthetic capabilities of microorganisms.

## RESULTS AND DISCUSSION

### Identification of novel RiPP precursor peptides

To investigate the diversity of YcaO-associated RiPP pathways, we focussed on standalone YcaO-domain proteins (i.e. those not fused to an additional domain) encoded in actinobacterial genomes, as the function of most of these standalone YcaO proteins are unknown.^[18]^ YcaO-domain proteins have been shown to catalyse various post-translational modifications, and so their precursor peptide substrates might greatly vary in amino acid sequence. 2,574 proteins were retrieved from GenBank, which were further filtered to 1,514 using a 95% maximum identity cut-off.^[20]^ Using these YcaO proteins as bait, RiPPER was used to retrieve associated short peptides and group them into families using similarity networking (40% minimum identity cut-off). This revealed a series of peptide families encoded within 8 kb of the *ycaO* genes (Figure S1, Supplementary Dataset 1). As expected, this included precursors to known RiPP families, including the bottromycins, thioviridamide-like molecules and thiopeptides (Figure 2). However, the most abundant peptide family consisted of 231 peptides whose RiPP products were completely unknown and only 78 were originally annotated as genes. Mapping these peptides to YcaO protein phylogeny revealed that this network belongs to a single clade of proteins, reflecting coevolution of YcaO proteins and their associated precursor peptides (Figure 3).

**Figure 2.**
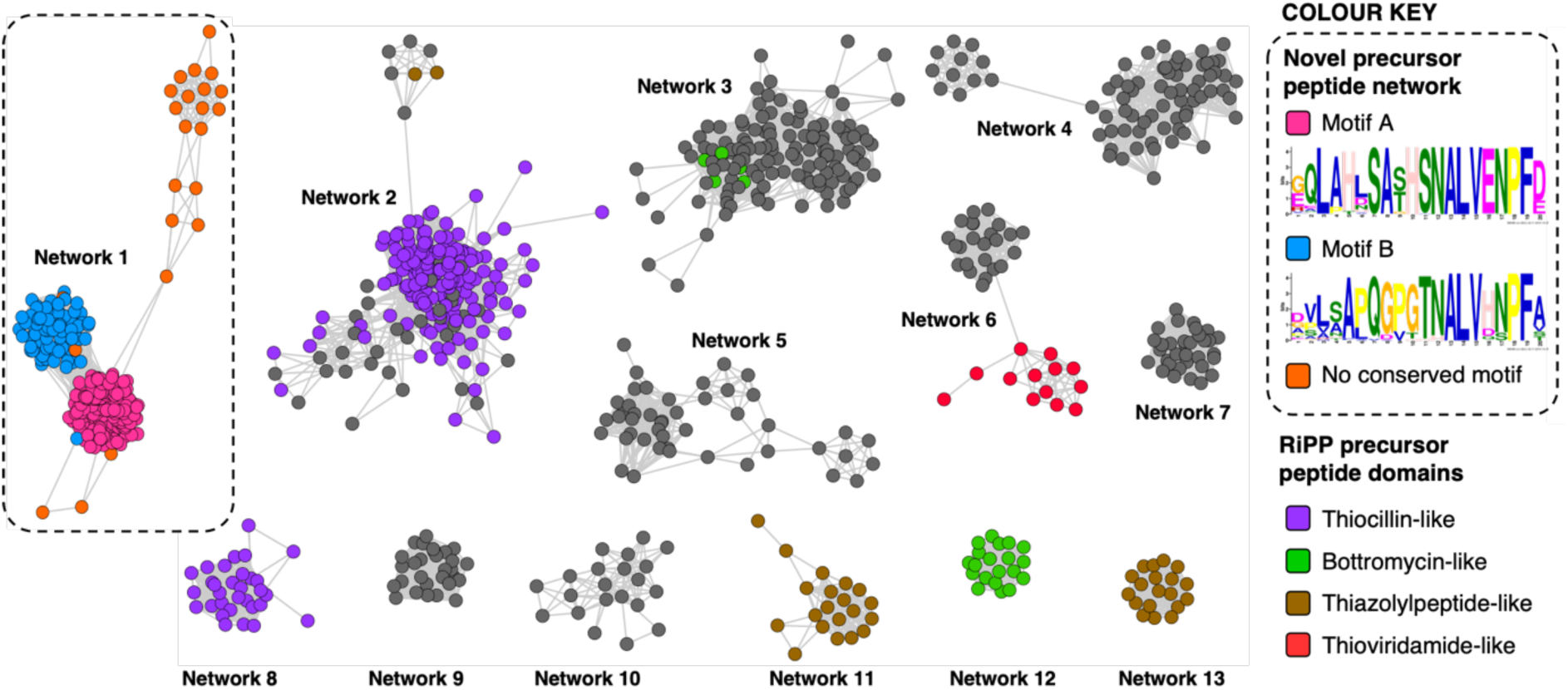
Sequence similarity networks of short peptides identified by RiPPER analysis of actinobacterial YcaO proteins. Peptides with homology to known RiPP classes are highlighted. Network 1 is the focus of this study, which is sub-divided based on the presence of different sequence motifs. A diagram of all identified precursor networks is shown in Figure S1.

**Figure 3.**
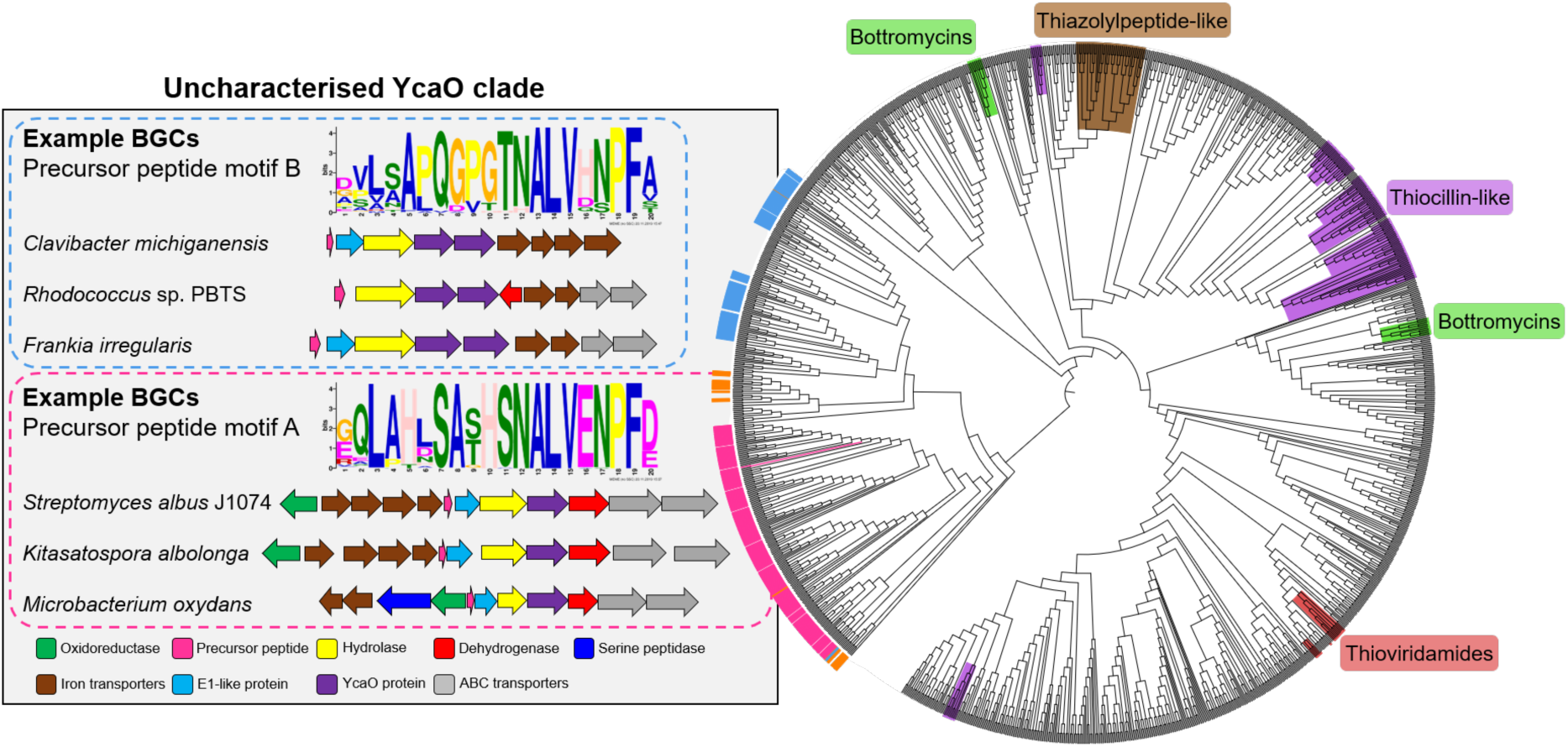
Phylogenetic tree of all standalone actinobacterial YcaO-domain proteins. The novel clades of YcaO-domain proteins identified from this analysis are highlighted according to their associated precursor peptide in pink (peptide with motif A), blue (motif B) and orange (no conserved motif), and examples of associated BGCs are shown. YcaO proteins associated with precursor peptides with known RiPP HMM domains from NCBI are also highlighted: red (thioviridamide family, NF033415), green (bottromycin family, NF033414), brown (thiazolylpeptide family, NF033400), and purple (thiocillin-like families, NF033482 and NF033401).

BLAST^[21]^ analysis of the YcaO proteins followed by RiPPER analysis revealed that six bacteria in the phylum Firmicutes also encoded related peptides near YcaO proteins. Overall, 237 novel precursor peptides were identified by RiPPER, and were present in eight orders, 22 bacterial families and 57 different genera (Supplementary Dataset 2). The identified precursor peptides varied in length between 31 and 89 residues, highlighting their diversity. A MEME analysis^[22]^ of the peptides identified two distinct sequence motifs (Figure 2), either of which appeared once, twice or three times in each precursor peptide. Whilst these motifs differ greatly in sequence, they both share a conserved ALV (alanine-leucine-valine) motif. In addition, 15 sequences lacked this motif and might therefore represent further precursor peptide diversification within this family. Notably, none of these peptides have serine/threonine/cysteine-rich regions that are characteristic of many precursors modified by YcaO proteins.^[18]^

To investigate the relationship between peptide sequence and YcaO-domain protein, these peptides were mapped to a phylogenetic tree of all actinobacterial standalone YcaO proteins (Figure 3). This clearly showed that this putative new family of precursor peptides is associated with a single clade of phylogenetically related YcaO-domain proteins. There are also different sub-clades that clearly associate with precursor peptides containing either motif A or motif B. A further similarity networking analysis of these 237 peptides using an 80% minimum identity cut-off resulted in a series of sub-families that mainly group by bacterial phylogeny (Figure S2), but there are some exceptions that might represent convergent evolution or horizontal transfer. These sub-families again map tightly to YcaO protein phylogeny (Figure S3).

### Genetic organisation of BGCs

The genes accompanying the YcaO and precursor peptide genes in this new family of RiPPs also show a high degree of conservation. MultiGeneBlast^[23]^ analysis of the newly identified BGCs revealed several subsets of BGCs whose genetic organisation correlates with the subclades identified within the family (Figures 3 and 4). The major one, found in over 90 BGCs (Figure S4A) contains the following set of conserved genes: four iron transporter genes with homology to the FecBCDE system (*amiF1-F4*), the putative precursor peptide (*amiA*), a conserved hypothetical protein (*amiB*), a hydrolase (*amiC*), the YcaO-domain protein (*amiD*), a flavin-dependent dehydrogenase (*amiE*) and two ABC transporters (*amiT1 and amiT2*). This subset of BGCs is usually associated with precursor peptides containing motif A (Figure 3). An exemplar of this BGC is found in the model streptomycete, *Streptomyces albus* J1074. This BGC also features a partially conserved hypothetical gene upstream of the iron transporters (*amiX*), which could also form part of the BGC. Other BGCs associated with the identified peptides have further diversity in their genetic composition. For example, many of the BGCs lack homologues of the *amiB*, *amiX* and *amiE* (dehydrogenase) genes, or contain additional hypothetical proteins with no identifiable conserved domains. (Figure S4B). Within these, a subset of BGCs found primarily in *Frankia*, *Rhodococcus* and *Clavibacter* contain two YcaO-domain proteins and are usually associated with precursor peptides containing motif B (Figure 3 and S4B).

**Figure 4.**
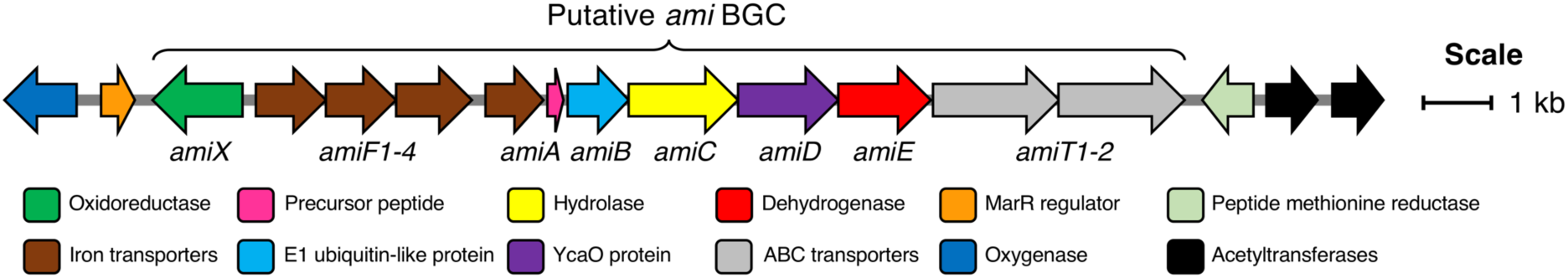
18.5 kb region of TAR cloned DNA from *S. albus* J1074 containing the putative *ami* BGC and additional flanking genes.

### Heterologous expression of the *S. albus* biosynthetic gene cluster

The BGC from *S. albus* J1074 was selected as a model for characterisation, as this contained the most widespread precursor peptide motif and BGC architecture. The resulting natural product would therefore represent the most abundant RiPP produced by the identified BGCs. We used transformation-associated recombination (TAR) cloning^[24]^ in yeast to capture an 18.5 kb region of genomic DNA from *S. albus* J1074 and generate plasmid pCAPSalbC. This contained the putative BGC (Figure 4), as well as additional genes upstream and downstream of this region that could feasibly have biosynthetic roles, including an oxygenase, a MarR transcriptional regulator, a peptide methionine sulfoxide reductase, and two acetyltransferases.

To determine the RiPP product of the BGC, an in-frame deletion of the precursor peptide gene, *amiA*, was generated in pCAPSalbC via PCR-targeting.^[25]^ “Wild type” pCAPSalbC and pCAPSalbC *ΔamiA* were introduced into *Streptomyces coelicolor* M1146, *Streptomyces lividans* and *Streptomyces laurentii* via intergeneric conjugation from *Escherichia coli*, and the resulting strains were fermented in multiple media. Untargeted metabolomic analysis of liquid chromatography–mass spectrometry (LC–MS) data revealed three major compounds (*m/z* 647.32, *m/z* 510.27 and *m/z* 409.22) that were produced by *S. coelicolor* M1146 containing the full cluster (*S. coelicolor* M1146-pCAPSalbC) but not the negative control strain that lacked the precursor peptide gene (Figure 5A).

**Figure 5.**
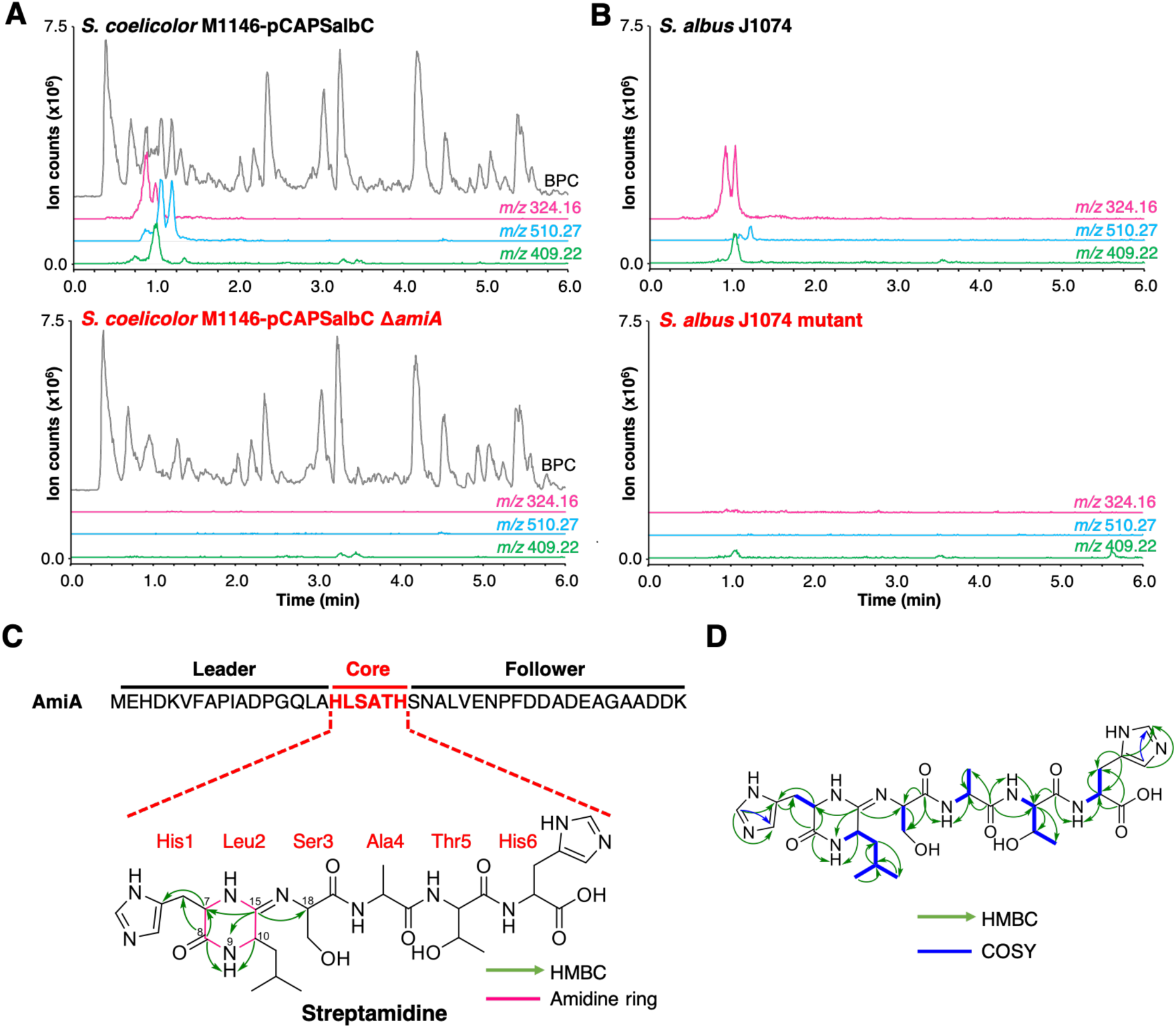
Discovery of streptamidine. A. LC-MS chromatograms of *S. coelicolor* M1146-pCAPSalbC compared with *S. coelicolor* M1146-pCAPSalbC Δ*amiA*. Extracted ion chromatograms of three BGC-associated compounds are shown: *m/z* 324.16 (pink), *m/z* 510.27 (blue) and *m/z* 409.22 (green). BPC = base peak chromatogram. B. Corresponding LC-MS data for *S. albus* J1074 compared with the *S. albus* J1074 BGC mutant. C. Precursor peptide AmiA with leader, core and follower regions highlighted along with the structure of streptamidine. This shows key HMBC correlations for the amidine ring. D. Detailed HMBC and COSY NMR correlation data for streptamidine (associated data shown in Figures S6-S14).

Based on similar tandem MS (MS^2^) fragmentation data (Figure S15), we proposed that these three compounds were related, and that the smaller masses might represent different intermediates or breakdown products of the final natural product, *m/z* 647.32 (also observed as [M+2H]^2+^, *m/z* 324.16). To further confirm that these compounds were being produced by the putative BGC, we grew the native strain, *S. albus* J1074, under the same conditions as for heterologous expression, and constructed a mutant disrupted in the YcaO gene (*amiD*) as a control. LC-MS analysis showed that all identified compounds (*m/z* 647.32, *m/z* 510.27 and *m/z* 409.22) were produced by *S. albus* J1074 but were not produced by the *ΔamiD* mutant (Figure 5B). These data provided further confirmation that these newly identified natural products were produced by the *ami* BGC in *S. albus* J1074.

### Structural elucidation of streptamidine

High-resolution LC-MS^2^ data indicated that the *m/z* 647.32 compound corresponds to the proton adduct of molecule with formula C_28_H_42_N_10_O_8_ (calculated [M+H]^+^ *m/z* 647.3260; observed *m/z* 647.3251). An analysis of all possible core peptides from AmiA along with a set of likely modifications indicated that the mass was consistent with a central HLSATH core region of the precursor peptide that had undergone dehydration. The formation of an oxazoline would be consistent with ATP-dependent cyclodehydration catalysed by the YcaO-domain protein. After large-scale fermentation, this compound was purified and the structure was elucidated by NMR (^1^H, ^13^C, COSY, HSQCed, HMBC, TOCSY and HSQC-TOCSY). This verified that the compound derived from the HLSATH core peptide. However, the chemical shifts for the side-chains of Ser3 (core peptide numbering) and Thr5 were consistent with unmodified amino acids rather than the corresponding heterocycles, whereas the ^13^C shift for the sp^2^ carbon between Leu2 and Ser3 (δ_C_ 157.1 ppm) differed from either an unmodified amide carbonyl or an oxazoline ring. Instead, HMBC correlations support a structure with a 6-membered amidine ring formed between the N-terminal amine and the carbonyl of Leu2. Correlations are shown in Figure 5 and include HMBC correlations between C15-H9 (δ_H_ 8.05), C15-H7 (δ_H_ 3.95-3.91) and C15-H18 (δ_H_ 4.30-4.27), which support the presence of an amidine ring. The chemical shift of C15 (δ_C_ 157.1 ppm) is similar to that of the corresponding carbons in the amidine rings of bottromycin (δ_C_ 157.9 ppm) in CDCl_3_^[26]^ and klebsazolicin (δ_C_ 156.8 ppm) in DMSO-d6.^[27]^

Due to the widespread presence of this BGC in streptomycetes and the rare amidine ring, this new compound was named streptamidine. The existence of multiple charge states and/or conformers may explain the peak shapes observed by LC-MS (Figure 5). High-resolution LC-MS^2^ analysis of the two other compounds identified (*m/z* 510.2668 and *m/z* 409.2195), indicated that these have masses that match those calculated for dehydrated HLSAT and HLSA peptides respectively (expected *m/z* 510.2671 and *m/z* 409.2196, respectively). Each compound produces a MS^2^ fragment consistent with the loss of a histidine moiety (obs. *m/z* 110.0723, exp. *m/z* 110.0713) and multiple other identical fragments (Figure S15). The prevalence of this BGC across Actinobacteria suggests an important function for streptamidine-like molecules. We hypothesised that this could be related to metal import, given the frequent association with *fecBCDE*-like genes. However, metal binding could not be detected with an iron-based CAS assay or with LC-MS-based binding assays with a range of metal ions [iron(II), cobalt(II), copper(II), magnesium(II), manganese(II), nickel(II) and zinc(II)]. Similarly, the streptamidine-null *S. albus ΔamiD* mutant was phenotypically identical to wild type *S. albus* under metal starvation conditions. No antibacterial or antifungal activity could be detected in assays against multiple indicator strains (Table S14).

### Identification of key biosynthetic machinery

To associate the production of the identified compounds to the cloned BGC, and to confirm a minimal set of genes required for streptamidine production, we generated a series of in-frame deletion mutants in the pCAPSalbC plasmid (Figure 6A). Deletion of *amiB* (hypothetical protein), *amiC* (hydrolase), *amiD* (YcaO-like protein), *amiE* (dehydrogenase), and *amiF1-F4* (iron transporters) abolished production of all pathway-associated compounds (Figure 6B and 6C). This suggested that these genes are essential for biosynthesis. In contrast, deletion of *amiT1-T2* (ABC transporters), *amiX* (putative oxidoreductase) gene, the MarR gene, the oxygenase gene, the peptide methionine sulfoxide reductase gene and the acetyltransferase genes had no effect on production of the compounds, indicating that these genes are not required for biosynthesis. This therefore enabled us to determine the minimal *ami* BGC (Figure 6A).

**Figure 6.**
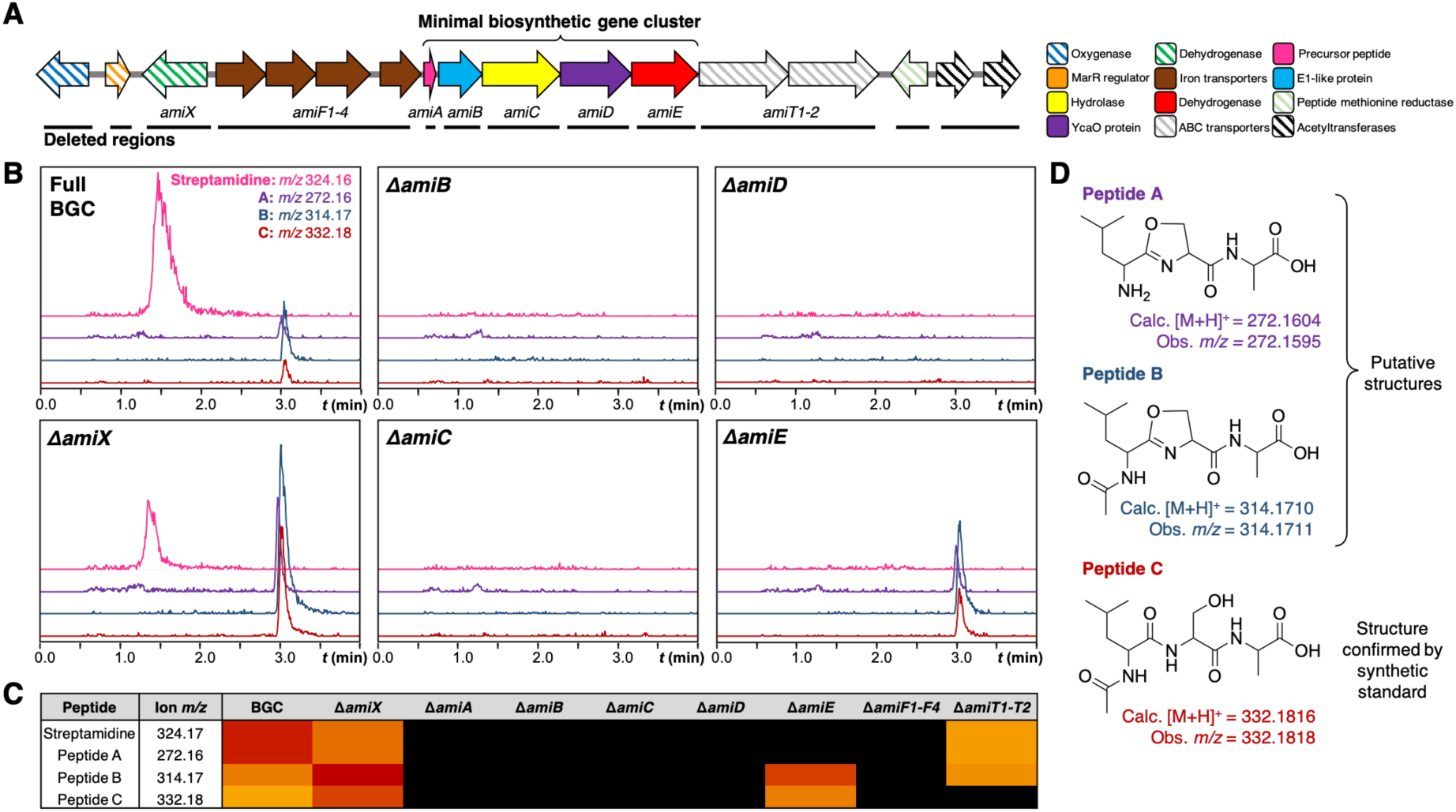
Mutational analysis of *ami* BGC. A. TAR cloned genetic region from *S. albus* with minimal biosynthetic gene cluster indicated. Striped arrows represent genes that are not essential for production of streptamidine, filled in arrows represent gene deletions that abolish production of streptamidine and are therefore essential for biosynthesis. The black lines beneath the genes indicate the regions that were independently deleted in this study. B. Metabolomic profiles of expressed gene cluster and key pathway mutants, including extracted ion chromatograms of shunt metabolite masses. C. Heat map indicating intensity of different metabolites produced by the gene cluster and pathway mutants. D. Predicted structures of streptamidine pathway shunt metabolites.

In addition to abolishing streptamidine production, the dehydrogenase mutant (*ΔamiE*) increased production of additional molecules: *m/z* 272.16, *m/z* 314.17 and *m/z* 332.18 (Figure 6B). The exact masses of these molecules indicated that these could derive from the Leu-Ser-Ala tripeptide within the core peptide (Figure 6D): dehydrated LSA (calc. *m/z* 272.1604, obs. *m/z* 272.1595), *N*-acetylated and dehydrated LSA (calc. *m/z* 314.1710, obs. *m/z* 314.1711) and *N*-acetylated LSA (calc. *m/z* 332.1816, obs. *m/z* 332.1818). This final molecule was confirmed by a comparison to a synthetic standard (Figure S17). Interestingly, production of these compounds was also increased in the oxidoreductase mutant (*ΔamiX*), although streptamidine was still produced by this strain (Figure 6B).

### Biosynthesis of the amidine ring

To date, only two RiPPs with amidine rings have been characterised: bottromycin and klebsazolicin.^[27, 28]^ In klebsazolicin biosynthesis, the BGC encodes one YcaO-domain protein that installs azole heterocycles and the amidine ring^[15]^ in cooperation with a partner E1-like protein and a dehydrogenase. The BGC for bottromycin encodes two YcaO proteins, where one is required for macroamidine formation^[29]^ and the other catalyses heterocyclisation of a cysteine residue to a thiazoline; both function without a partner protein.^[14, 30]^ In the case of streptamidine, the gene deletion data are consistent with a role in cyclisation for AmiB and the YcaO protein, AmiD. Conventional sequence analysis did not identify any conserved domains for AmiB, but structural modelling with Phyre2^[31]^ predicts that it has a homologous structure to residues 4-315 of the cyanobactin heterocyclase TruD,^[32]^ encompassing a RiPP recognition element^[33]^ and an E1-like domain. This suggests that AmiB and AmiD cooperate to catalyse cyclisation in an analogous way to heterocycle-forming YcaO proteins.

Deletion of genes encoding dehydrogenase AmiE and hypothetical protein AmiX led to the accumulation of molecules with accurate masses consistent with dehydrated Leu-Ser-Ala derived from the core peptide. This hints that an oxazoline-containing intermediate could be formed before the final amidine-containing structure is generated, and that the dehydrogenase is somehow important for cyclisation. In klebsazolicin biosynthesis, Travin *et al.*^[15]^ proposed that an intermediate ring structure might form on the Ser3 residue before the amidine is ultimately produced, which could potentially happen in streptamidine biosynthesis (Figure S18). In relation to this mechanism, the serine in position 3 of the streptamidine core peptide is conserved across motif A-containing precursor peptides (Figure S19). Deletion of the hydrolase gene *amiC*, which is predicted to remove the leader peptide prior to amidine formation, abolishes streptamidine production, as a free *N*-terminal amine on His1 is required for cyclisation.

It is somewhat unexpected that the dehydrogenase AmiE is essential for streptamidine production despite the lack of an oxidation in streptamidine. Possible explanations include: (a) AmiE is fulfilling a key structural role for proper cyclisation activity; (b) AmiE is catalytic but a reductase reverses this activity; (c) the oxidised part of the peptide is hydrolysed from the streptamidine core region. AmiE is highly dissimilar to characterised azole-forming dehydrogenases (~10% identity to the microcin B17 dehydrogenase McbC) but Phyre2^[31]^ analysis indicates that it has a HY motif that structurally aligns with the catalytic KY residues of the microcin B17 dehydrogenase McbC^[34]^ (Figure S21). Detailed biochemical experiments will be required to determine the true role of the dehydrogenase and order of biosynthetic steps, but the production of high levels of streptamidine by both the heterologous host and wild type *S. albus* J1074 indicates that it is a major product of the pathway.

## CONCLUSIONS

This work shows that the development and application of targeted genome mining tools is a valuable approach to identify uncharacterised novel biosynthetic gene clusters. In this study we identified over 230 novel BGCs that are widespread in well-studied bacteria such as *Streptomyces*, as well as understudied genera such as *Frankia* and *Rhodococcus.* Guided by genetic and metabolomic analyses, we isolated and characterised streptamidine, a previously overlooked amidine-containing RiPP from a model streptomycete. This represents a very rare example of an amidine-containing peptide in nature, yet our analysis indicates that these compounds could be widespread.

The *S. albus* J1074 precursor peptide sequence contains motif A identified by MEME analysis. These precursor peptides are encoded in BGCs with very similar genetic architectures, which suggests that a range of homologues of streptamidine may be produced in nature. An analysis of mass spectral databases using MASST (Mass Spectrometry Search Tool)^[35]^ identified a molecule with identical mass and MS^2^ fragmentation to streptamidine in a marine actinomycete MS dataset (Figure S22, MassIVE MSV000078679). However, the precursor sequences containing motif B feature very distinct amino acid sequences and lie within varied BGC architectures (Figure S4B). These BGCs might therefore collectively produce a wide range of structurally distinct RiPPs. This highlights that there is still a vast amount of untapped chemical diversity to be discovered from uncharacterised RiPP BGCs, as have other recent studies that have used genomics-led approaches to identify widespread novel RiPP chemistry.^[7,36,37]^

The biological role of streptamidine remains unknown, but the prevalence and widespread distribution of streptamidine-like BGCs in nature suggests a beneficial role for the producing organism. This is comparable to other widespread natural products whose activities remain a mystery.^[38, 39]^ The resulting molecules may therefore have an important role that could be linked to signalling or development rather than inhibitory activity, which warrants further investigations into this new family of RiPP. Interestingly, Metelev *et al* observed that the six-membered amidine ring of klebsazolicin is essential for its unique ability to form a compact conformation inside the ribosome exit tunnel, which is more obstructive than other ribosome inhibitors such as macrolides. It was therefore suggested that amidine-containing structures could be viable starting points for the development of bioactive molecules such as antibiotics, with the amidine cycle acting as part of a functional ‘warhead’.^[27]^ Therefore, the discovery of a widespread family of amidine-containing molecules represents an exciting opportunity to explore this possibility. Furthermore, the extent of YcaO proteins that are associated with uncharacterised precursor peptide families (Figure 2, Supporting Dataset 1) highlights the wealth of RiPP diversity that remains to be discovered.

## MATERIALS AND METHODS

### Strains and culture conditions

All strains, plasmids, culture media and primers used in this work are described in the supplementary methods (Tables S1, S2, S3, S4, S5, S6). Unless otherwise specified, all *Streptomyces* strains were grown at 28 °C on solid SFM for spore growth, solid SFM supplemented with 10 mM MgCl_2_ for conjugations, liquid TSB for seed cultures and liquid SM12 media for fermentations. Other media used during screening trials include liquid R5, BPM and SM14. Liquid cultures were grown with shaking at 250 rpm. Spores and mycelium stocks were kept at −20 °C in 20% glycerol. *Saccharomyces cerevisiae* VL6–48N^[40]^ was used for transformation-associated recombination (TAR) cloning and was grown at 30 °C with shaking at 250 rpm in YPD medium. Recombinant yeast selection was performed using selective media SD+CSM-Trp complemented with 5-fluoorotic acid (Fluorochem, 1 mg mL^-1^). *Escherichia coli* DH5α was used for transformation and propagation of DNA plasmids. For gene deletions, *E. coli* DH5α BT340 was used for Flp-FRT recombination and *E. coli* BW25113/pIJ790 was used for Lambda-Red mediated recombination^[25]^. pIJ790-carrying strains were grown at 30 °C for plasmid replication, and Flp-FRT recombination was performed at 42 °C. *E. coli* ET12567/pR9604 and *E. coli* ET12567/pUZ8002 were used to transfer DNA to *Streptomyces* by intergeneric conjugation. All *E. coli* strains were grown in LB medium at 37 °C unless otherwise specified. *E. coli* hygromycin selection was performed in solid DNA media. *E. coli* cell stocks were kept at −20 °C in 20% glycerol. Antibiotic selection was carried out using the following final concentrations of antibiotic: kanamycin 50 μg mL^-1^, apramycin 50 μg mL^-1^, hygromycin 50 μg mL^-1^, nalidixic acid 25 μg mL^-1^, chloramphenicol 25 μg mL^-1^ and carbenicillin 100 μg mL^-1^.

### Analysis of YcaO-domain proteins

All actinobacterial standalone YcaO-domain proteins were identified in NCBI Genbank using CDART (Conserved Domain Architecture Retrieval Tool)^[41]^ (2,574 proteins). These were filtered using EFI-EST^[20]^ to 2,338 sequences after excluding proteins smaller than 350 AA, and further filtered to 1,514 proteins using a 95% identity cut-off. Corresponding accession numbers were submitted to Batch Entrez and the resulting sequence files aligned using MUSCLE.^[42]^ This alignment was used to construct a maximum likelihood tree using RAxML-HPC2 on XSEDE with default settings on the CIPRES Science Gateway (https://www.phylo.org/). The tree was visualised with the interactive Tree Of Life (iTOL).^[43]^

### Retrieval and analysis of precursor peptides and BGCs

The 1,514 YcaO-domain protein accessions were used as the input for RiPPER^[7]^ (https://github.com/streptomyces/ripper) with default settings. Some of the precursor peptide sequences obtained from RiPPER analysis were duplicated due to the presence of more than one YcaO protein in some BGCs. 29 duplicated precursor sequences were manually removed prior to subsequent analysis. Peptide similarity networking of the precursor peptide sequences were created using EGN (Evolutionary Gene and genome Network)^[44]^ and visualised with Cytoscape 2.8.3.^[45]^ Multiple sequence alignments of precursor peptides were performed using ClustalW^[46]^ via MEGA7^[47]^ using default settings, and motifs were searched for using the MEME tool^[48]^ in the MEME suite (http://meme-suite.org/index.html) using classic mode with the site distribution as Any Number of Repetitions and searching for 3 motifs. The captured genomic regions were visualised and analysed in Artemis^[49]^ and VectorNTI,^[50]^ and putative BGCs were compared using MultiGeneBlast^[23]^ (http://multigeneblast.sourceforge.net/). Conserved protein domains were analysed using NCBI conserved domain search (https://www.ncbi.nlm.nih.gov/Structure/cdd/wrpsb.cgi) and Phyre2.^[31]^

### TAR cloning and heterologous expression of *Streptomyces albus* J1074 gene cluster

A vector to capture the gene cluster from *S. albus* J1074 genomic DNA (gDNA) was constructed by Gibson assembly between a linearised pCAP03 vector^[51]^ and two single-strand oligonucleotides (Salb_TAR_fw and Salb_TAR_rv). The forward and reverse oligonucleotides had 34 and 36 nucleotide homology sequences with pCAP03 respectively. These were designed to generate a vector with 50 and 49 nucleotide homology sequences with upstream and downstream regions of the gene cluster respectively, either side of an AvrII restriction site. pCAP03 was digested with XhoI and NdeI, and 100 ng linearised plasmid and 10 pmol of each oligonucleotide were incubated with 5 µL ligase-free Gibson assembly reaction (100 mM Tris-HCl pH 7.5, 10 mM MgCl_2_, 0.2 mM each dNTPs, 10 mM DTT, 1 mM NAD, 5% PEG-8000, 0.1125 units T5 exonuclease, 0.375 units Phusion polymerase, 10 µL total reaction volume) and incubated at 50 °C for 2 hours in a BioRad T100 ThermoCycler. 10 μL assembly reaction was then transformed into *E. coli* DH5α by chemical transformation selected with kanamycin. Colonies containing the correct capture vector were identified by PCR using primers pCAP_sp and pCAP_asp, and the plasmid was isolated using a Promega Wizard Plus SV Minipreps DNA Purification System.

Genomic DNA from *S. albus* J1074 was digested with NsiI and SmlI, and the capture vector was linearised between the capture arms with AvrII. Digested material was transformed into *S. cerevisiae* VL6–48N by spheroplast polyethylene glycol 8000 transformation. Successful gene cluster capture by pCAP03 was confirmed by colony PCR. For yeast-colony PCR, each colony was resuspended in 50 μL 1 M sorbitol (Fisher) and 2 μL of zymolyase (5 U μL^-1^) added to each cell suspension and incubated at 30 °C for 1 hour. Cell suspensions were then boiled for 10 min, centrifuged (15 s, 1,000 × g) and 1 μL of the supernatant was analysed by PCR using primers Salb_TARscr_Fw and Salb_TARscr_Rv. The plasmids from four positive clones were recovered and transformed into electrocompetent *E. coli* DH5α for further analytical digest of the purified construct with HindIII-HF and SrfI. *E. coli* ET12567/pR9604 was transformed with pCAPSalbC by electroporation, and transformants were then used to transfer pCAPSalbC into *S. coelicolor* M1146 by intergeneric conjugation. Nalidixic acid and kanamycin-resistant exconjugants containing integrated pCAPSalbC (*S. coelicolor* M1146-pCAPSalbC) were verified by PCR using Promega GoTaq polymerase with primers Salb_TARscr_Fw and Salb_TARscr_Rv.

### Construction of *S. albus* J1074 pathway mutant

A fragment of DNA corresponding to the translationally coupled YcaO and hydrolase genes was PCR cloned with restriction sites for EcoRI and HindIII. The DNA fragment was digested with EcoRI and HindIII and ligated into the pKC1132 plasmid digested with EcoRI and HindIII. The resulting DNA construct was isolated and transferred into *S. albus* J1074 via intergenic conjugation with *E. coli* ET12567/pUZ8002 selected with apramycin, chloramphenicol and kanamycin. Exconjugants resistant to apramycin were validated by PCR to confirm that the YcaO and hydrolase genes had been disrupted.

### Fermentation and metabolite screening by LC-MS

Seed cultures of *S. coelicolor* M1146-pCAPSalbC were prepared by fermentation in a 50 mL flask containing 5 mL of TSB with kanamycin selection for 48 h. 500 μL seed culture was used to inoculate 10 mL SM12, SM14, BPM and R5 in 50 mL Falcon tubes with caps replaced by foam bungs. Control strains carrying the TAR clone with a precursor peptide gene deletion were cultured in the same way for comparison. Seed cultures of *S. albus* J1074 were grown in the same way with no antibiotic selection, and the *S. albus ΔamiD* mutant seed cultures were grown with apramycin selection. All fermentations were conducted in triplicate and incubated at 28 °C with shaking at 230 rpm. 1 mL culture samples were taken at day 4, mixed with one volume of methanol and agitated for 30 min at room temperature. These mixtures were then centrifuged (15,871 × g, 5.5 min) and 800 μL of the resulting supernatant was transferred to glass vials for liquid chromatography–mass spectrometry (LC–MS) analysis. LC-MS samples were analysed on a Shimadzu Nexera X2 UHPLC coupled to a Shimadzu IT-TOF mass spectrometer. 5 μL samples were injected onto a Phenomenex Luna Omega 1.6-µm Polar C18 column (50 mm by 2.1 mm, 100 Å) set at a temperature of 40 °C and eluting with a linear gradient of 0–60% methanol in water + 0.1% formic acid over 6 minutes with a flow-rate of 0.6 mL min^-1^. Positive mode mass spectrometry data was collected between *m/z* 200 and 2,000. Untargeted comparative metabolomics was carried out on data from triplicate samples using Profiling Solution 1.1 (Shimadzu) with an ion *m/z* tolerance of 100 mDa, a retention time tolerance of 0.1 min and an ion intensity threshold of 70,000 units.

For accurate mass analysis, mass spectra were acquired by LC–MS on a Synapt G2-Si mass spectrometer equipped with an Acquity UPLC (Waters). Samples were injected onto an Acquity UPLC BEH C18 column, 1.7 μm, 1 × 100 mm (Waters) and eluted with a gradient of (B) acetonitrile/0.1% formic acid in (A) water/0.1% formic acid with a flow rate of 0.08 ml min^-1^ at 45 °C. The concentration of B was kept at 1% for 1 min followed by a gradient up to 60% B over 10 min, and up to 99% over 2 min MS data were collected with the following parameters: resolution mode, positive ion mode, scan time 0.5 s, mass range *m/z* 50–1200 (calibrated with sodium formate), capillary voltage = 3.0 kV; cone voltage = 40 V; source temperature = 110 °C; desolvation temperature = 250 °C. Leu-enkephalin was used to generate a lock-mass calibration with *m/z* = 556.2766 measured every 30 s during the run.

Streptamidine-like molecules were searched for using a MASST search^[35]^ at Global Natural Products Social Molecular Networking (https://gnps.ucsd.edu/ProteoSAFe/static/gnps-splash.jsp) with the following settings: parent mass tolerance = 2 Da; min matched peaks = 5; ion tolerance = 0.5 Da; score threshold = 0.7; library = speclib.

### Deletion of genes in *Streptomyces albus* J1074 biosynthetic gene cluster

Mutations of the *S. albus* J1074 BGC were carried out using an *E. coli*-based Lambda-Red-mediated PCR-targeting strategy,^[25]^ which allowed substitution of genes in pCAPSalbC by a PCR-generated cassette containing the apramycin resistance gene *aac(3)-IV*. Resistance cassettes were amplified by PCR using a pIJ773-derived cassette lacking OriT as a template, which allowed the elimination of the apramycin resistance cassette after Flp-FRT recombination in *E. coli* DH5α BT340, to create mutants with an in-frame 81 bp scar in the place of the original gene sequence. The PCR-targeting mutant versions of pCAPSalbC were introduced into *S. coelicolor* M1146 by *E. coli* ET12567/pR9604-mediated intergeneric conjugation and selected by resistance to nalidixic acid and kanamycin.

### Complementation of deleted genes

Constructs for the complementation of mutants were obtained by high-fidelity PCR amplification (Q5 polymerase) of each of these genes, digestion of the PCR product with NdeI and HindIII and cloning by ligation (T4 DNA ligase, Invitrogen) into pIJ10257 digested with NdeI and HindIII.^[52]^ Ligation mixtures were transformed into chemically competent *E. coli* DH5α and the plasmids were recovered by miniprep and then sequenced. The constructs were introduced into the corresponding *S. coelicolor* M1146-pCAPSalbC mutants by *E. coli* ET12567/pR9604-mediated intergeneric conjugation. Exconjugants were selected by resistance to nalidixic acid, kanamycin and hygromycin.

### Purification of streptamidine

Four 2-litre flasks containing 0.5 L of SM12 were each inoculated with 25 mL of *S. coelicolor* M1146-pCAPSalbC TSB seed culture grown for 48 hours at 28 °C. After four days fermentation at 28 °C with shaking at 250 rpm, the cultures were centrifuged to remove debris, combined and filtered to yield approximately 1.5 L of crude extract. The crude extract was extracted with ethyl acetate (3 × 1.5 litres). The aqueous layer was further extracted with 1-butanol (3 × 1.0 litres). The resulting aqueous extract was concentrated to 50 mL using a Buchi rotary evaporator and subjected to solid-phase chromatography (SPE) on a HP20 cartridge using a gradient of H2O-MeOH (100:0 to 0:100). Methanol and water were removed using a Buchi rotary evaporator until the samples were concentrated to approximately 10 mL. Samples were then subject to semi-preparative HPLC using a Phenomenex Luna PFP(2) column (5 μm, 250 x 10 mm) with a gradient of aqueous 0.1% formic acid-MeOH (98:2 to 90:10) over 35 minutes with a flow rate of 2 mL min^-1^. The compound was monitored at a UV wavelength of 210 nm and fractions were assessed by LC-MS. Fractions containing streptamidine were combined and freeze dried. A final purification step was then carried out using a semi-preparative Luna Omega Polar C18 column (5 μm, 250 x 10 mm), with an isocratic gradient of aqueous 0.1% formic acid-MeOH (90:10) for 16 minutes followed by a wash gradient from 90:10 to 5:95 over 5 minutes with a flow rate of 2.8 mL min^-1^. The compound was monitored at a UV wavelength of 210 nm and fractions assessed by LC-MS.

### Structural elucidation of streptamidine

Pure streptamidine (1.4 mg) was dissolved in 600 μL DMSO-d6 from an individual vial and subjected to a series of 1D and 2D nuclear magnetic resonance (NMR) experiments on a Bruker Ascend 600 MHz instrument at 298 K. The NMR experiments carried out were Proton (64 scans), Carbon (25,000 scans), HSQCed (100 scans), HMBC (64 scans), COSY (16 scans), TOCSY (32 scans) and HSQC-TOCSY (64 scans). Spectra were analysed using Bruker TopSpin 3.5 and Mestrelab Research Mnova 14.0 software. NMR data are reported in Figures S5-S14 and Table S9.

### Antimicrobial assays

10 mL cultures of each indicator strain (Table S14) were grown in LB (YPD for *Candida utilis*) overnight at 37 °C (30 °C for *C. utilis*). 100 µL of each culture was then used to inoculate a 10 mL subculture of each strain in the same medium, which were grown for 5 hours at 37 °C (30 °C for *C. utilis*). 1 mL of each culture was then mixed with 14 mL molten LB agar, which was poured into plates. Once solidified, three 1 cm diameter plugs were taken from each agar plate, which were then separately loaded with 50 µL streptamidine (1 mg mL^-1^), 50 µL kanamycin, apramycin or nalidixic acid (1 mg mL^-1^) as a positive control and 50 µL water as a solvent control. Plates were incubated overnight at 37 °C (30 °C for *C. utilis*).

### Metal binding assays

For the CAS assay, 500 μL of CAS assay solution (prepared as described by Alexander and Zuberer^[53]^) was mixed with 10 µL increasing concentrations of streptamidine from 1.5 µM to 25 µM. For LC-MS binding assays, solutions of 10 mM metal salts were prepared (FeCl_3_, CoCl_2_, CuCl_2_, MgCl_2_, MnSO_4_, NiSO_4_, ZnCl_2_, dissolved in 10 mM HCl) and 500 μL of each were mixed with 20 µL streptamidine (15 µM). 5 μL samples were analysed on a Shimadzu Nexera X2 UHPLC coupled to a Shimadzu IT-TOF mass spectrometer with a Phenomenex Luna Omega 1.6-µm Polar C18 column (50 mm by 2.1 mm, 100 Å) set at a temperature of 40 °C and eluting with a linear gradient of 0–60% methanol in water + 0.1% formic acid over 6 minutes with a flow-rate of 0.6 mL min^-1^. Positive mode mass spectrometry data was collected between *m/z* 200 and 2,000.

For metal starvation experiments, a minimal medium was prepared (2 g K_2_SO_4_, 3 g K_2_HPO_4_, 1 g NaCl, 5 g NH4Cl, 0.005 mg CuSO_4_, 0.035 mg MnSO_4_.H_2_O, 2 mg ZnSO_4_.7H_2_O, 80 mg MgSO_4_.7H_2_O, 100 mg CaCl_2_.2H_2_O, 2.5% glycerol) as described in Muller and Raymond^[54]^ in glassware washed with EDTA. A series of the same minimal media omitting either copper, zinc or iron were also prepared. Seed cultures of *S. albus* J1074 and the *S. albus ΔamiD* mutant were prepared by fermentation in a 50 mL flask containing 5 mL of TSB (with apramycin selection for the *ΔamiD* mutant) for 48 h. 500 μL each seed culture was used to inoculate 10 mL minimal media and each metal dropout media in 50 mL Falcon tubes with caps replaced by foam bungs. Cultures were grown for 4 days and 1 mL samples were taken for LC-MS analysis as described above.

## ACKNOWLEDGEMENTS

This work was funded by a Biotechnology and Biological Sciences Research Council (BBSRC) Norwich Research Park Doctoral Training Partnership grant (BB/M011216/1) for A.H.R., a Royal Society University Research Fellowship (A.W.T.), and BBSRC MET and MfN Institute Strategic Programme grants (BB/J004596/1 and BBS/E/J/000PR9790) for the John Innes Centre (JIC). We are very grateful for the technical assistance at JIC provided by Lionel Hill, Gerhard Saalbach and Carlo de Oliveira Martins for LC-MS, Martin Rejzek for HPLC and Sergey Nepogodiev for NMR. We thank Vladimir Larionov (National Cancer Institute, NIH, USA) for *S. cerevisiae* VL6-48N and Bradley Moore (Scripps Institution of Oceanography, University of California San Diego, USA) for pCAP03. We are thankful to Barrie Wilkinson, Tom Eyles and Javier Santos-Aberturas for helpful discussions.

