## Supplementary Information for "Discovery and characterisation of an amidine-containing ribosomally-synthesised peptide that is widely distributed in nature"

**Table S1** Strains used in this study.

| Strain | Genotype/description | Application |
|---|---|---|
| *Saccharomyces cerevisiae* VL6-48N[1] | *MATα*, *his3-Δ1, trp1-Δ1, ura3-Δ1, lys2, ade2-101, met14 cir$^{o}$* | TAR cloning |
| *Escherichia coli* DH5α | F$^{-}$ ϕ80 *lacZ*ΔM15 Δ(*lacZYA-argF*)U169 *recA*1 *endA*1 *hsdR*17 ($r_k^-$, $m_k^+$) *phoAsupE*44 *thi*-1 *gyrA*96 *relA*1 λ$^{-}$ | Transformation and maintenance of plasmids and constructs |
| *E. coli* ET12567/pUZ8002 | *dam-13::Tn9 dcm-6 hsdM* Cml$^{R}$, carrying helper plasmid pUZ8002 | Conjugations with pKC1132 disruption construct |
| *E. coli* ET12567/pR9604 | *dam-13::Tn9 dcm-6 hsdM* Cml$^{R}$, carrying helper plasmid pR9604 | Conjugations with TAR cloned BGC, mutated BGC and complementation constructs |
| *E. coli* DH5α /BT340 | *E. coli* DH5α carrying BT340 plasmid. | Gene deletions |
| *E. coli* BW25113/pIJ790 | *(Δ(araD-araB)567, ΔlacZ4787(::rrnB-4), lacIp-4000(lacIQ), λ-,rpoS369(Am), rph-1, Δ(rhaD-rhaB)568, hsdR514. Plasmid: pIJ790 [oriR101], [repA101(ts)], araBp-gam-be-exo* | Gene deletions |
| *Streptomyces albus* J1074[2] | Restriction-defective derivative ($R^-M^-$) of *S. albus* G | Genetic source of BGC |
| *Streptomyces coelicolor* M1146[3] | Δ*act* Δ*red* Δ*cpk* Δ*cda* | Heterologous expression of gene cluster |

**Table S2** Plasmids used in study.

| Plasmid | Features | Resistance marker | Application |
|---|---|---|---|
| pCAP03[4] | ARSH4/CEN6-Trp1, pUC ori, C31 int-attP-oriT-aph, URA3, ADH1 | Kanamycin | TAR cloning |
| pKC1132[5] | Conjugative vector, non-integrative, lacZa | Apramycin | *S. albus* pathway disruption |
| pIJ773[6] | oriT, non-conjugative, flippase recognition target (FRT) sites | Apramycin | Gene deletions via PCR targeting |
| pIJ10257[7] | ΦBT1, p*erm*E* | Hygromycin | Genetic complementation of mutants |

**Table S3** Constructs generated in study.

| Construct | Resistance marker | Application |
|---|---|---|
| pCAPSalbC | Kanamycin | pCAP03-based plasmid containing the TAR cloned *S. albus* BGC |
| pCAPSalbC_ΔPP | Kanamycin | pCAPSalbC with deletion of precursor peptide gene |
| pCAPSalbC_ΔYcaO | Kanamycin | pCAPSalbC with deletion of YcaO gene |
| pCAPSalbC_ΔE1 | Kanamycin | pCAPSalbC with deletion of E1-like gene |
| pCAPSalbC_ΔDehy | Kanamycin | pCAPSalbC with deletion of dehydrogenase gene |
| pCAPSalbC_ΔHydro | Kanamycin | pCAPSalbC with deletion of hydrolase gene |
| pCAPSalbC_ΔOxido | Kanamycin | pCAPSalbC with deletion of oxidoreductase gene |
| pCAPSalbC_ΔABC | Kanamycin | pCAPSalbC with deletion of set of ABC transporter genes |
| pCAPSalbC_ΔIrTr | Kanamycin | pCAPSalbC with deletion of set of iron transporter genes |
| pCAPSalbC_ΔPepMet | Kanamycin | pCAPSalbC with deletion of peptidyl methionine gene |
| pCAPSalbC_ΔOxyg | Kanamycin | pCAPSalbC with deletion of oxygenase gene |
| pCAPSalbC_ΔAcet | Kanamycin | pCAPSalbC with deletion of both acetyltransferase genes |
| pKC1132_SalbYH | Apramycin | Construct for disruption of *S. albus* BGC in native host |
| pIJSalb_YcaO | Hygromycin | Construct for complementation of YcaO deletion |
| pIJSalb_E1 | Hygromycin | Construct for complementation of E1 deletion |
| pIJSalb_Dehy | Hygromycin | Construct for complementation of dehydrogenase deletion |

**Table S4** Media used in study.

| Medium | Application | Ingredients per 1 L, made up with milliQ water |
|---|---|---|
| LB | *E. coli* | 10 g Bacto-tryptone, 5 g yeast extract, 10 g NaCl, adjust to pH 7 with NaOH |
| DNA | *E. coli* | 4 g Difco Nutrient Broth powder, 10 g agar |
| TSB | *Streptomyces* | 17 g tryptone, 3 g phytone, 5 g NaCl, 2.5 g $K_2HPO_4$, 2.5 g glucose |
| SFM | *Streptomyces* | 20 g mannitol, 20 g soya flour, 100 mM $CaCl_2$, 20 g agar |
| YPD | *Saccharomyces* | 10 g yeast extract, 20 g peptone, 20 g glucose (15 g agar), 0.004% adenine |
| SD-Trp | *Saccharomyces* | 5 g $(NH_4)_2SO_4$, 1.7 g YNB-AA, 20 g glucose, 0.74 g CSM-Trp (20 g agar), 0.004% adenine |
| Top selective agar | *Saccharomyces* | 182 g sorbitol, 22 g dextrose, 30 g agar, 0.0002% 5-FOA, 0.004% adenine |
| Bottom selective agar | *Saccharomyces* | 182 g sorbitol, 22 g dextrose, 20 g agar, 0.0002% 5-FOA, 0.004% adenine |
| R5 | *Streptomyces* | 103 g sucrose, 0.25 g $K_2SO_4$, 10.12 g $MgCl_2.6H_2O$, 10 g glucose, 0.1 g casamino acids, 2 mL trace element solution, 5 g yeast extract, 5.73 g TES buffer |
| SM12 | *Streptomyces* | 10 g soy flour, 50 g glucose, 4 g peptone, 4 g beef extract, 1 g yeast extract, 2.5 g NaCl, 5 g $CaCo_3$, adjust to pH 7.6 with KOH |
| SM14 | *Streptomyces* | 10 g glucose, 20 g soy peptone, 5 g meat extract, 5 g NaCl, 0.01 g $ZnSO_4.7H_2O$, adjust to pH 7.0 with KOH |
| BPM | *Streptomyces* | 15 g starch, 5 g yeast extract, 10 g soy flour, 5 g NaCl, 3 g $CaCO_3$, 25 µg/mL $CoCl_2$ |

**Table S5** Solutions used for TAR cloning.

| Solution | Ingredients per 100 mL, made up with milliQ water |
|---|---|
| 10x nitrogen bases | 1.9 g YNB-AA, 1.9 g CSM-Trp, 5 g $NH_4SO_4$ |
| 100x adenine | 1 g adenine, 74 mM HCl |
| SPE | 10 mM HEPES buffer pH 7.5, 100 mM EDTA pH 8, 18.2 g sorbitol |
| SOS | 15 mM $CaCl_2$, 0.25 g yeast extract, 18.2 g sorbitol, 1 g peptone |
| STC | 10 mM Tris.HCl pH 7.5, 10 mM $CaCl_2$, 18.2 g sorbitol |
| PEG | 10 mM Tris.HCl pH 7.5, 10 mM $CaCl_2$, 20 g PEG8000 |

### Primers

All primers were ordered from Eurofins Genomics and purified by HPSF. Short primers were ordered with a synthesis scale of 0.01 µmol and long primers for PCR targeting were ordered with a synthesis scale of 0.05 µmol.

**Table S6** Primers used in study.

| Primer name | Sequence (5'-3') | Use | Restriction site |
|---|---|---|---|
| SalbCap_Fw | GCTGCCGGGCCGGCTCCTAGGTCTACATCGGGG ACATCAGCGACGCCCGTCCCGCGAGTCTTCCGAT GCCGTTAATTAAGCCACTATTTATACCATGGGAG GCGTCAAAC | Construction of pCAP03-derived capture vector for TAR cloning | - |
| SalbCap_Rv | TGTCCCCGATGTAGACCTAGGAGCCGGCCCGGC AGCTGACGGGTCAGCCACGGCAGGAACCGCGG GCCGTCATATGTCGAAAGCTACATATAAGGAACG TGCTG | | - |
| Salb_ClusScr_Fw | GCAGGACGGAACCGAGGGATG | Screening for cluster capture | - |
| Salb_ClusScr_Rv | TGGGAGAGGATCGCCTCGGC | | - |
| Salb_YHMut_Fw | GATACAAAGCTTGACTGGATACGCGCCCAGC | Amplification of DNA fragment for pathway disruption in *S. albus* | HindIII |
| Salb_YHMut_Rv | GATACAGAATTCGCTCACCTCCAGGCCGGACC | | EcoRI |
| Salb_TAR_PPDel_Fw | CGTCCACCACGCATCGAACTGAATGGAGCTCAAC T CATGATTCCGGGGATCCGTCGACC | Precursor peptide gene deletion | - |
| Salb_TAR_PPDel_Rv | TGTCAGCCGGCCGCCGTCACCGGCGGCCTGGGC TGACTATGTAGGCTGGAGCTGCTTC | | - |
| Salb_TAR_E1Del_Fw | GCCCATCCCCTCGTACTCCATCCGACGGAGGTTT CCGTGATTCCGGGGATCC | E1-like gene deletion | - |
| Salb_TAR_E1Del_Rv | CGGGCCGCTCCCCCTTGCGGGCAGCGGGGCCA CCGGCGGTGTAGGCTGGAGCTGCTTC | | - |
| Salb_TAR_YcaODel_Fw | GACCTGCCGATGACCGCCGCCCTGCCCCTCGAC GCCCTCATTCCGGGGATCCGTCGACC | YcaO gene deletion | - |
| Salb__TAR_YcaODel_Rv | GGGAGCCGGGGGTCACATGTGCGGGGCGGGGG CGAGGGTTGTAGGCTGGAGCTGCTTC | | - |
| Salb_TAR_HydrDel_Fw | GCTGCCCGCAAGGGGGAGCGGCCCGGCATGAC GCCCGCGATTCCGGGGATCCGTCGACC | Hydrolase gene deletion | - |
| Salb_TAR_HydrDel_Rv | GGGCGTCGAGGGGCAGGGCGGCGGTCATCGGC AGGTCTCTGTAGGCTGGAGCTGCTTC | | - |
| Salb_TAR_DehyDel_Fw | CGCCCCCGCCCCGCACATGTGACCCCCGGCTCC CCCATGATTCCGGGGATC | Dehydrogenase gene deletion | - |
| Salb_TAR_DehyDel_Rv | GGGTGAGGTGGTCGGGGGCGGGCCGTGCGCGG CGGCTCATGTAGGCTGGA | | - |
| Salb_TAR_OxidoDel_Fw | CGCCCGGCCCCCGCACCCCCTACCGAGGAGTTC CCCGTGATTCCGGGGATCCGTCGACC | Oxidoreductase gene deletion | - |
| Salb_TAR_OxidoDel_Rv | GCGACACCCTGGCCCGCGCCTGGCCGAGCTGAG GAGTCATGTAGGCTGGAGCTGCTTC | | - |

| Primer name | Sequence (5'-3') | Use | Restriction site |
|---|---|---|---|
| Salb_TAR_MarRDel_Fw | AGGGACGCTACACGACGAGCGAGGAGACCCGCG ACCATGATTCCGGGGATCCGTCGACC | MarR regulator gene deletion | - |
| Salb_TAR_MarRDel_Rv | ACGGGGCGGAGGCGGACCCGGTGGGGCGGTGA CTCCTCATGTAGGCTGGAGCTGCTTC | | - |
| Salb_TAR_Oxyg_Fw | TCGAAGTTCACCATCCAGCAGCGCGCGGTTCCC GCGATGATTCCGGGGATCCGTCGACC | Oxygenase gene deletion | - |
| Salb_TAR_Oxyg_Rv | CGTTCTCGCTCATGCGCGTCTCCTTCCTCGGCTC GTTCATGTAGGCTGGAGCTGCTTC | | - |
| Salb_TAR_IrTr_Fw | ACGCGTCCCCGCAGGACGGAACCGAGGGATGAA GCCATGATTCCGGGGATCCGTCGACC | Iron transporters gene deletions | - |
| Salb_TAR_IrTr_Rv | GGTCGGTTCGTCGAGGAGGAGGGTCCGGGTGTC CTGGGCTGTAGGCTGGAGCTGCTTC | | - |
| Salb_TAR_ABC_Fw | CGACCACCTCACCCCGCCCGGAGAGACGTACCC GCGATGATTCCGGGGATCCGTCGACC | ABC transporters gene deletions | - |
| Salb_TAR_ABC_Rv | CGTGTGCGTGCGCGTGTACGCACGCTTCGGTGC GGGTCATGTAGGCTGGAGCTGCTTC | | - |
| Salb_TAR_PepMet_Fw | CCGGGCGCATGTCGATGCCAGTCGGGAGCACAG CGTATGATTCCGGGGATCCGTCGACC | Peptide methionine sulfoxide reductase MsrA gene deletion | - |
| Salb_TAR_PepMet_Rv | CCCCGGCTCCTCGGTCCGGTGAAGGAGTGCTGT GGCTCATGTAGGCTGGAGCTGCTTC | | - |
| Salb_TAR_Acet_Fw | CCGCCGGTCCGCACGCCACCGGGAGGGGCCCA CCGCATGATTCCGGGGATCCGTCGACC | Acetyl transferase and maltose-O-acetyltransferase gene deletions | - |
| Salb_TAR_Acet_Rv | GGGCCCGGCCCCCTTCGCGTGTACGTACGGGCC CCGTCATGTAGGCTGGAGCTGCTTC | | - |
| SalbCycl_PE_ Fw | GATACACATATGACCAGCAGCCGACTCGCC | Complementation of E1-like protein | NdeI |
| SalbCycl_PE_ Rv | GATACAAAGCTTGTGGTCGCGGGCGTCATGC | | HindIII |
| SalbYcaO_PE_Fw | GATACACATATGACCGCCGCCCTGCCC | Complementation of YcaO | NdeI |
| SalbYcaO_PE _Rv | GATACAAAGCTTGGGGAGCCGGGGGTCACATG | | HindIII |
| SalbDehy_PE _Fw | GATACACATATGACCCCTGACGCCACCCTCG | Complementation of dehydrogenase | NdeI |
| SalbDehy_PE _Rv | GATACAAAGCTTGCTCATCGGGCGGCTCCCAG | | HindIII |
| PPDel_screen_Fw | GCGGCTGGCCGGTCTGTTAC | Screening precursor peptide gene deletion | - |
| PPDel_screen_Rv | CGGCTGCTGGTCACGGAAACC | | - |
| CyclDel_screen_Fw | GGTGCCGCGGACGACAAGTAG | Screening E1-like gene deletion | - |
| CyclDel_screen_Rv | GTCGTACGGGGTGCGGATCAG | | - |
| YcaODel_screen_Fw | ACCAGGCTGCGCGTCGAGA | Screening YcaO gene deletion | - |
| YcaODel_screen_Rv | AGGTGGTCGAGGTCGACGGG | | - |
| HydrDel_screen_Fw | CTCCTCACCGCCGACCTCCTC | Screening hydrolase gene deletion | - |
| HydrDel_screen_Rv | GTGCCGAGCGAGACCCGGT | | - |
| DehyDel_screen_Fw | TCGACCTGACCACCGAGGACG | Screening dehydrogenase gene deletion | - |
| DehyDel_screen_Rv | ACCAGGGCGAGCAGGGCG | | - |
| OxidoDel_Screen_Fw | CATCCCTCGGTTCCGTCCTG | | - |

| Primer name | Sequence (5’-3’) | Use | Restriction site |
|---|---|---|---|
| OxidoDel_Screen_Rv | AGCACCCTGATCCGGCTGAC | Screening oxidoreductase gene deletion | - |
| Salb_oxyg_scr_Fw | CAGTTGAGGGGCGGATCGTTC | Screening oxygenase deletion | - |
| Salb_oxyg_scr_Rv | GAAAGGCCCAGCTGGGCGTC | | - |
| Salb_IronTr_scr_Fw | CAGAGGCGTCCCACGCGTC | Screening iron transporters deletion | - |
| Salb_IronTr_scr_Rv | GAACGGCGTGGCGACTGCC | | - |
| Salb_ABCTr_scr_Fw | CTCACCCCGCCCGGAGAGAC | Screening ABC transporters deletion | - |
| Salb_ABCTr_scr_Rv | GTGCGTGCGCGTGTACGCAC | | - |
| Salb_PepMet_scr_Fw | GACCCCGGCTCCTCGGTCC | Screening peptide methionine sulfoxide reductase MsrA deletion | - |
| Salb_PepMet_scr_Rv | CGCATGTCGATGCCAGTCGG | | - |
| Salb_acet_scr_Fw | GTGACACCAAGGTGCCGCGAAC | Screening acetyl transferase and maltose-O-acetyltransferase deletions | - |
| Salb_acet_scr_Rv | GGCCCCCTTCGCGTGTACG | | - |
| MarR_screen_Fw | TCAGCCCGACCGGTCCTG | Screening MarR deletion | - |
| MarR_screen_Rv | CGACCACGCCGAGGAGGTC | | - |
| pIJ10257_Fw | TTCGAGTGGCGGCTTGCG | Screening for pIJ10257 insert | - |
| pIJ10257_Rv | CAAACGGCATTGAGCGTCAGC | Screening for pIJ10257 insert | - |

**Table S7** Accurate masses of streptamidine and related compounds.

| Description | Formula | Calc. $[M+H]^+$ | Obs. *m/z* | Error (ppm) |
|---|---|---|---|---|
| Streptamidine (modified HLSATH) | $C_{28}H_{42}N_{10}O_8$ | 324.1666 ($[M+2H]^{2+}$) | 324.1666 | 0.00 |
| | | 647.3260 | 647.3251 | 1.39 |
| Predicted modified HLSAT | $C_{22}H_{35}N_7O_7$ | 510.2671 | 510.2668 | 0.59 |
| Predicted modified HLSA | $C_{18}H_{28}N_6O_5$ | 409.2194 | 409.2195 | -0.24 |
| Predicted dehydrated LSA | $C_{12}H_{21}N_3O_4$ | 272.1604 | 272.1595 | 3.31 |
| Predicted acetylated and dehydrated LSA | $C_{14}H_{23}N_3O_5$ | 314.1710 | 314.1711 | -0.32 |
| Predicted acetylated LSA | $C_{14}H_{25}N_3O_6$ | 332.1816 | 332.1818 | -0.60 |

**Table S8** Proteins encoded in the streptamidine biosynthetic gene cluster.

| Name | Accession | Pfam domain | Predicted function/domain | Size (AA) |
|---|---|---|---|---|
| AmiX | WP_085479279.1 | No Pfam match | Oxidoreductase | 417 |
| AmiF1 | WP_085479278.1 | PF01497 | Iron transporter complex | 329 |
| AmiF2 | WP_085479277.1 | PF01032 | Iron transporter complex | 344 |
| AmiF3 | WP_085479276.1 | PF01032 | Iron transporter complex | 351 |
| AmiF4 | WP_0854792751 | PF00005 | Iron transporter complex | 278 |
| AmiA | AMM12575.1 | No Pfam match | Precursor peptide | 44 |
| AmiB | WP_085479274.1 | No Pfam match | E1-like protein | 281 |
| AmiC | WP_129865485.1 | PF02129 | Hydrolase | 508 |
| AmiD | WP_008409979.1 | PF02624 | YcaO-domain | 455 |
| AmiE | WP_129857084.1 | No Pfam match | Dehydrogenase | 435 |
| AmiT1 | WP_049977232.1 | PF00005 | ABC transporter | 581 |
| AmiT2 | WP_049977233.1 | PF00005 | ABC transporter | 594 |

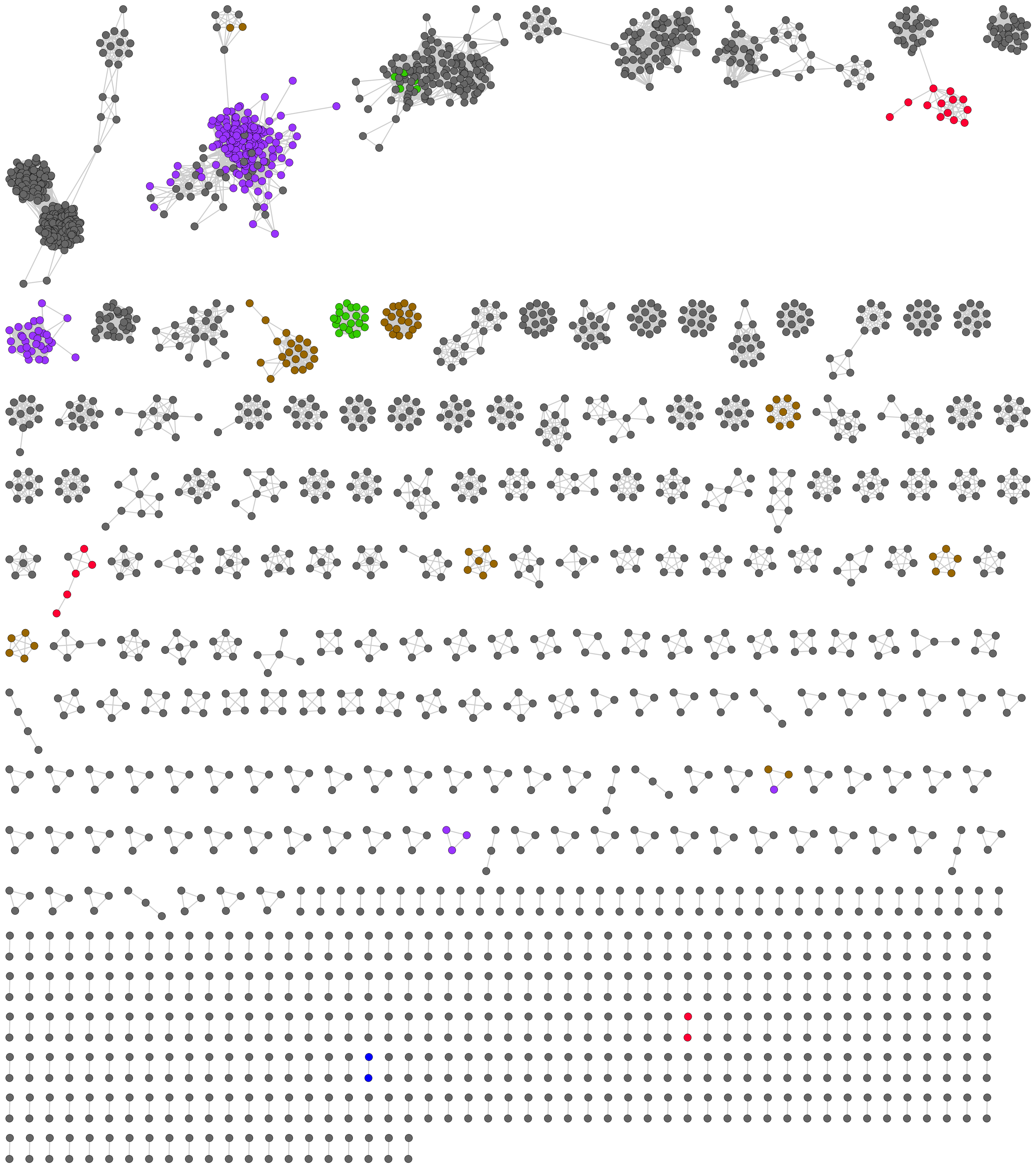

**Figure S1** Precursor peptide networks associated with Actinobacterial YcaO proteins. Precursors with homology to known RiPPs are highlighted: green = bottromycin family (NCBI HMM domain NF033414), brown = thiazolylpeptide families (NF033400 and NF033399), red = thioviridamide family (NF033415), purple = thiocillin families (NF033482 and NF033401), dark blue = lasso peptide family (NF033521).

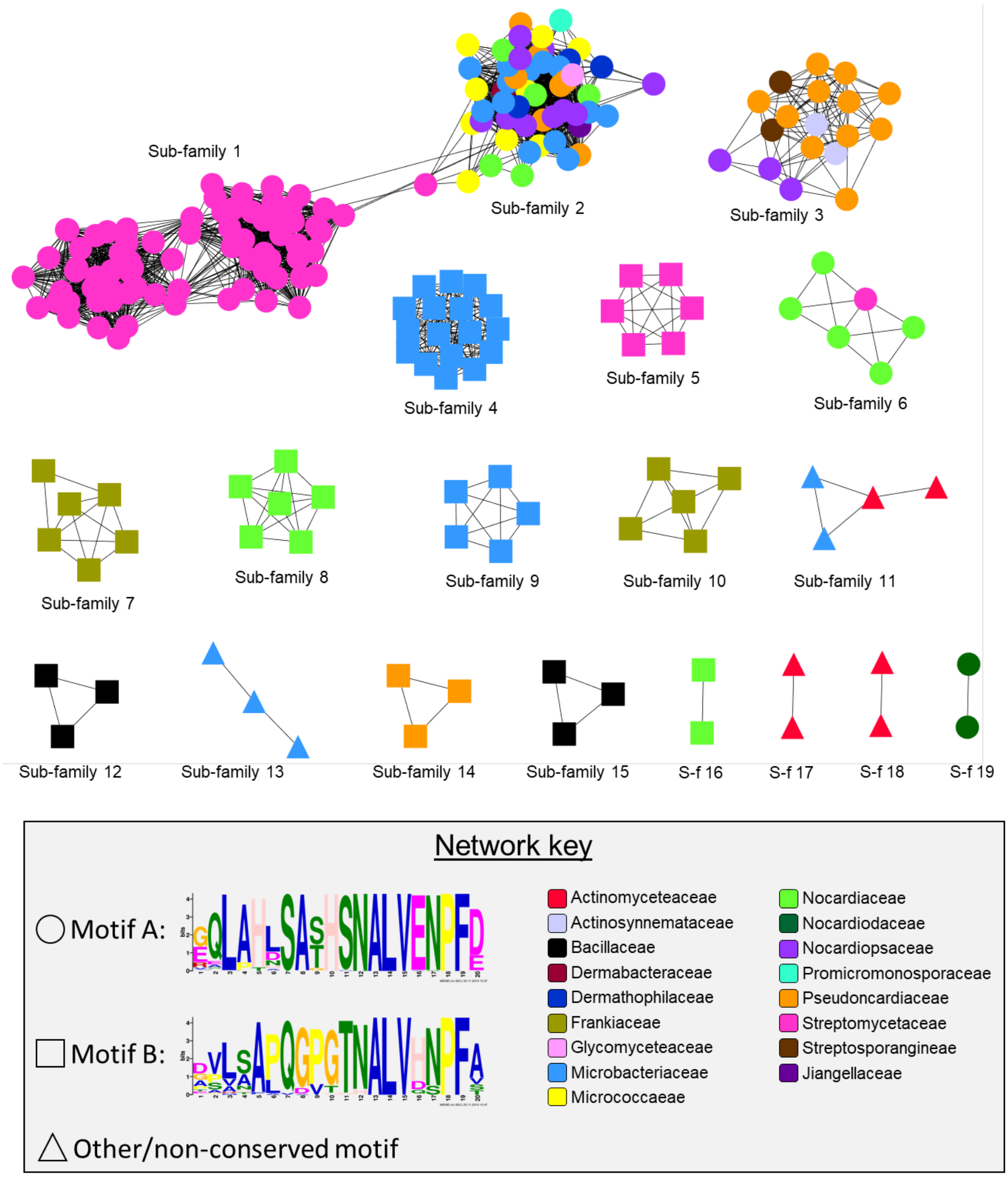

**Figure S2** EGN (Evolutionary Gene and Genome Network)[8] analysis of the 237 peptide sequences identified with RiPPER[9] (network one from Figure 1 and Figure S1) using an identity cut-off of 80%. Similar sequences are clustered together in sub-families and nodes are colour coded by the associated bacterial family. Peptides containing motif A are indicated with a circular node and peptides containing motif B are indicated by a square node. Precursors containing neither motif are indicated by a triangular node. Motifs A and B were identified by MEME[10] analysis of the 237 sequences. Each motif appears once, twice or three times in 208 of the precursor sequences.

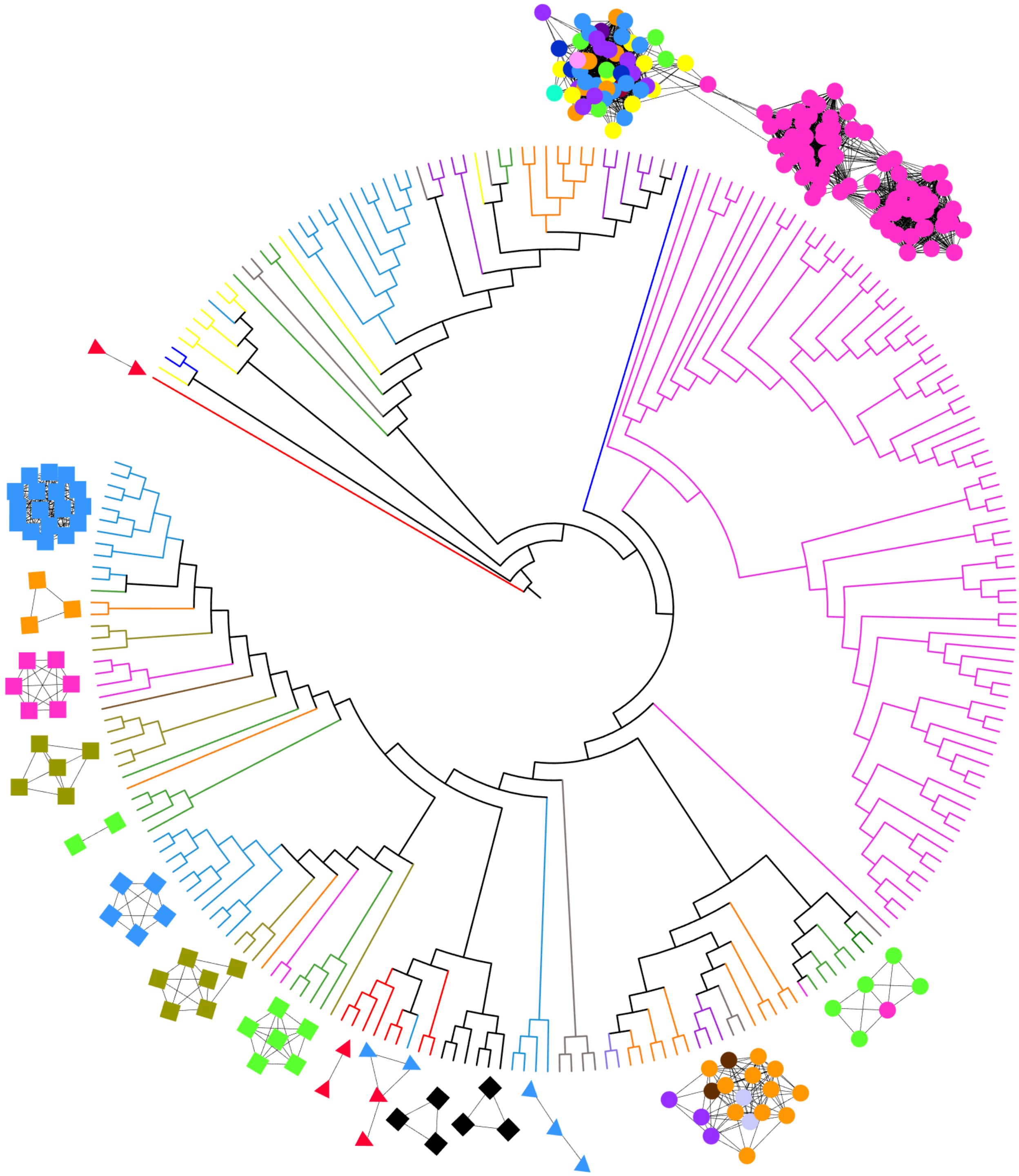

**Figure S3** Phylogenetic tree of YcaO proteins associated with network 1 of putative precursor peptides. The associated networks of precursor peptide sub-families are mapped (80% identity networks from Figure S2). Nodes and sub-families are colour-coded by bacterial family, showing that precursor peptides appear to have co-evolved with their cognate YcaO (colour coding as in Figure S2). YcaO sequence alignment and phylogenetic tree created as described in main paper. Tree visualised using iTOL.[11]

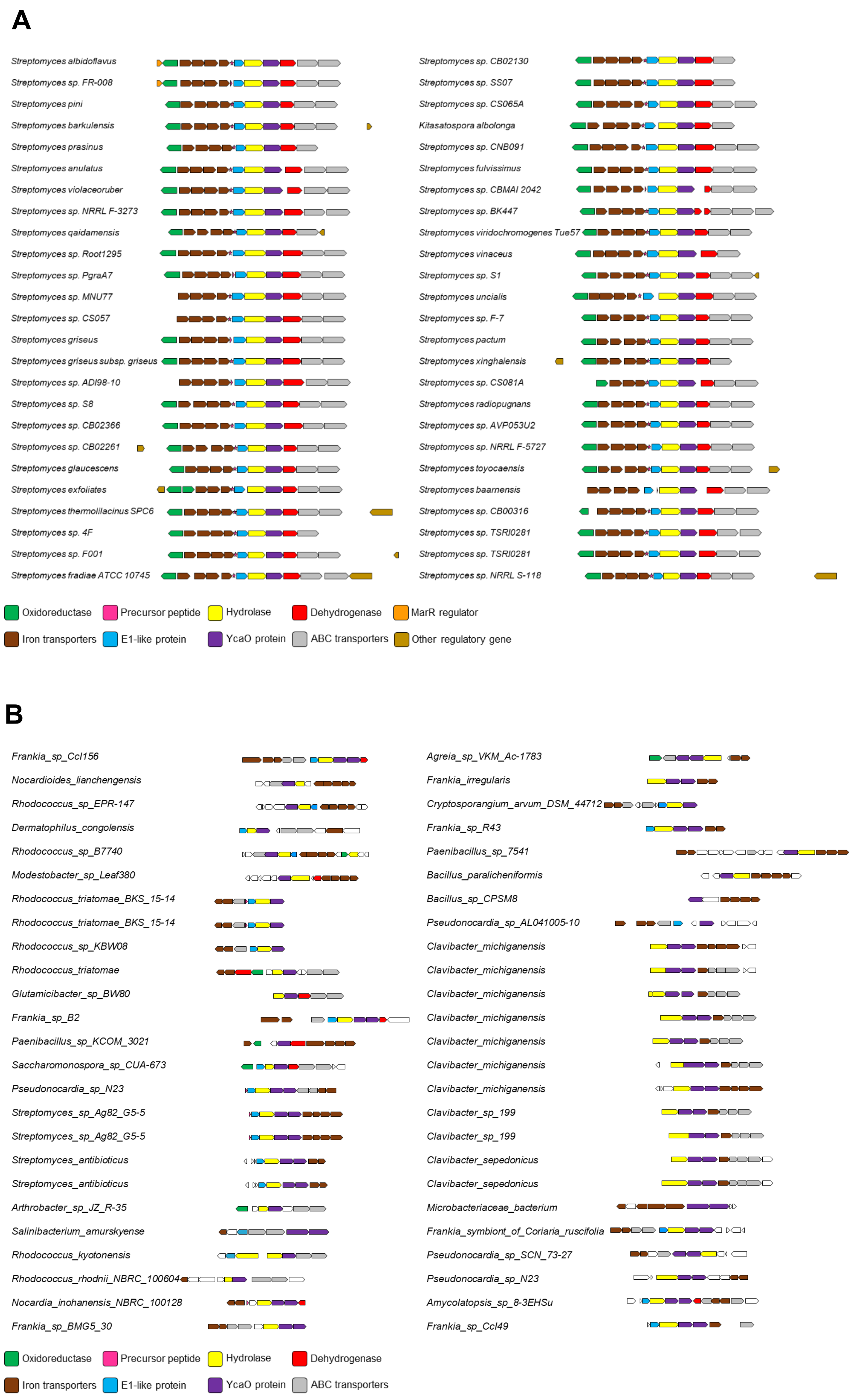
A
Streptomyces albidoflavus
Streptomyces sp. FR-008
Streptomyces pini
Streptomyces barkulensis
Streptomyces prasinus
Streptomyces anulatus
Streptomyces violaceoruber
Streptomyces sp. NRRL F-3273
Streptomyces qaidamensis
Streptomyces sp. Root1295
Streptomyces sp. PgraA7
Streptomyces sp. MNU77
Streptomyces sp. CS057
Streptomyces griseus
Streptomyces griseus subsp. griseus
Streptomyces sp. ADI98-10
Streptomyces sp. S8
Streptomyces sp. CB02366
Streptomyces sp. CB02261
Streptomyces glaucescens
Streptomyces exfoliates
Streptomyces thermolilacinus SPC6
Streptomyces sp. 4F
Streptomyces sp. F001
Streptomyces fradiae ATCC 10745
Streptomyces sp. CB02130
Streptomyces sp. SS07
Streptomyces sp. CS065A
Kitasatospora albolonga
Streptomyces sp. CNB091
Streptomyces fulvissimus
Streptomyces sp. CBMAI 2042
Streptomyces sp. BK447
Streptomyces viridochromogenes Tue57
Streptomyces vinaceus
Streptomyces sp. S1
Streptomyces uncialis
Streptomyces sp. F-7
Streptomyces pactum
Streptomyces xinghaiensis
Streptomyces sp. CS081A
Streptomyces radiopugnans
Streptomyces sp. AVP053U2
Streptomyces sp. NRRL F-5727
Streptomyces toyocaensis
Streptomyces baarnensis
Streptomyces sp. CB00316
Streptomyces sp. TSRI0281
Streptomyces sp. TSRI0281
Streptomyces sp. NRRL S-118
Oxidoreductase
Precursor peptide
Hydrolase
Dehydrogenase
MarR regulator
Iron transporters
E1-like protein
YcaO protein
ABC transporters
Other regulatory gene
B
Frankia_sp_CcI156
Nocardioides_lianchengensis
Rhodococcus_sp_EPR-147
Dermatophilus_congolensis
Rhodococcus_sp_B7740
Modestobacter_sp_Leaf380
Rhodococcus_triatomae_BKS_15-14
Rhodococcus_triatomae_BKS_15-14
Rhodococcus_sp_KBW08
Rhodococcus_triatomae
Glutamicibacter_sp_BW80
Frankia_sp_B2
Paenibacillus_sp_KCOM_3021
Saccharomonospora_sp_CUA-673
Pseudonocardia_sp_N23
Streptomyces_sp_Ag82_G5-5
Streptomyces_sp_Ag82_G5-5
Streptomyces_antibioticus
Streptomyces_antibioticus
Arthrobacter_sp_JZ_R-35
Salinibacterium_amurskyense
Rhodococcus_kyotonensis
Rhodococcus_rhodnii_NBRC_100604
Nocardia_inohanensis_NBRC_100128
Frankia_sp_BMG5_30
Agreia_sp_VKM_Ac-1783
Frankia_irregularis
Cryptosporangium_arvum_DSM_44712
Frankia_sp_R43
Paenibacillus_sp_7541
Bacillus_paralicheniformis
Bacillus_sp_CPSM8
Pseudonocardia_sp_AL041005-10
Clavibacter_michiganensis
Clavibacter_michiganensis
Clavibacter_michiganensis
Clavibacter_michiganensis
Clavibacter_michiganensis
Clavibacter_michiganensis
Clavibacter_michiganensis
Clavibacter_sp_199
Clavibacter_sp_199
Clavibacter_sepedonicus
Clavibacter_sepedonicus
Microbacteriaceae_bacterium
Frankia_symbiont_of_Coriaria_ruscifolia
Pseudonocardia_sp_SCN_73-27
Pseudonocardia_sp_N23
Amycolatopsis_sp_8-3EHSu
Frankia_sp_CcI49
Oxidoreductase
Precursor peptide
Hydrolase
Dehydrogenase
Iron transporters
E1-like protein
YcaO protein
ABC transporters

**Figure S4 (previous page)** A. First 50 examples of homologous gene clusters identified by MultiGeneBlast[12] analysis of the *S. albus* BGC. The core set of putative biosynthetic genes are present in all examples, which are predominantly from the *Streptomyces* genus and are associated with precursor peptides containing motif A. Surrounding genes that do not have predicted biosynthetic function within the RiPP biosynthetic pathway have been omitted, with the exception of genes annotated as having a regulatory function. B. BGCs 151-200 identified from MultiGeneBlast analysis of the *S. albus* BGC. These BGC examples are distinct from the first 50 examples, as many encode two YcaO proteins, additional hypothetical proteins (in white), and generally lack the *amiE*, *amiB* and *amiX* genes. Additionally, this subset of BGCs is usually associated with precursor peptides containing motif B.

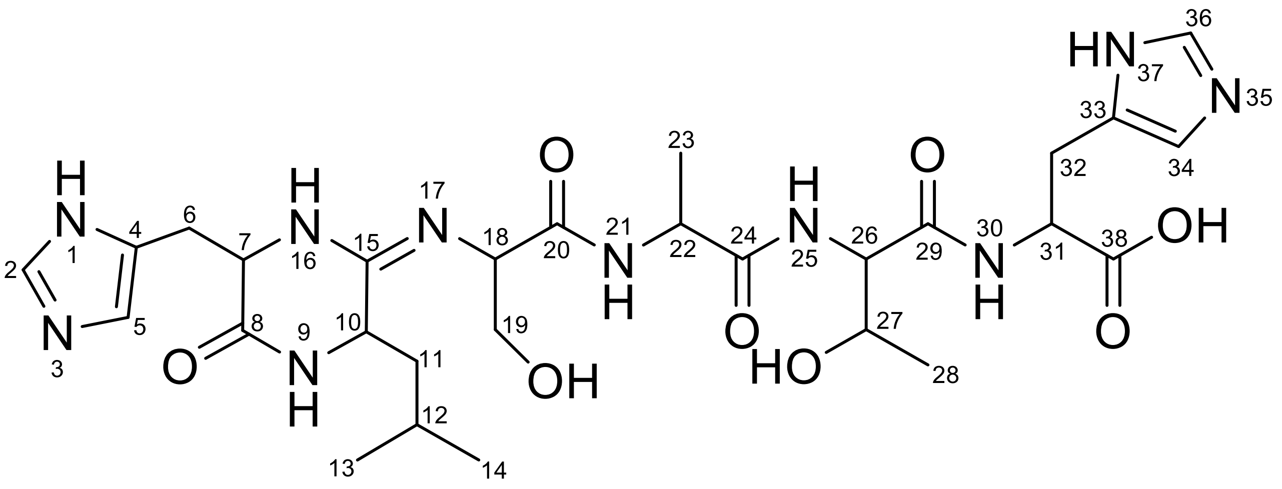

**Figure S5** Structure of streptamidine with numbered atoms.

**Table S9** NMR chemical shift assignments for streptamidine in DMSO-$d_6$

| C/N number | Amino acid | $\delta_C$ (ppm) | $\delta_H$ (ppm) | Multiplicity | Coupling constant (Hz) |
|---|---|---|---|---|---|
| 1 | His1 | -- | -- | -- | |
| 2 | His1 | 134.7 | 7.48 | s | |
| 3 | His1 | -- | | | |
| 4 | His1 | 131.6[a] | -- | -- | |
| 5 | His1 | 121.0[b] | 6.67 | s | |
| 6 | His1 | 32.6 | 2.84<br>2.75 | dd<br>dd | $J_{6a,6b}$ = 14.1, $J_{6a,7}$ = 5.2<br>$J_{6a,6b}$ = 14.1, $J_{6b,7}$ = 7.4 |
| 7 | His1 | 58.8 | 3.95-3.91 | m | |
| 8 | His1 | 170.7 | -- | -- | |
| 9 | Leu2 | -- | 8.05 | s | |
| 10 | Leu2 | 51.0 | 3.82 | d | $J_{10,11a}$ = 10.8 |
| 11 | Leu2 | 45.9 | 1.41-1.35, 0.77-0.71 | m, m | |
| 12 | Leu2 | 23.8 | 1.64-1.57 | m | |
| 13 | Leu2 | 21.4 | 0.82 | d | $J_{12,13}$ = 6.4 |
| 14 | Leu2 | 24.1 | 0.78 | d | $J_{12,14}$ = 6.6 |
| 15 | Leu2 | 157.1 | -- | -- | |
| 16 | His1 | -- | | | |
| 17 | Ser3 | -- | -- | -- | |
| 18 | Ser3 | 57.1 | 4.30-4.27 | m | |
| 19 | Ser3 | 62.2 | 3.62 | d | $J_{18,19}$ = 5.9 |
| 20 | Ser3 | 171.4 | -- | -- | |
| 21 | Ala4 | -- | 8.09 | d | $J_{21,22}$ = 7.0 |
| 22 | Ala4 | 48.9 | 4.41-4.35 | m | |
| 23 | Ala4 | 18.7 | 1.25 | d | $J_{22,23}$ = 7.1 |
| 24 | Ala4 | 172.9 | -- | -- | |
| 25 | Thr5 | -- | 7.88 | d | $J_{25,26}$ = 8.6 |
| 26 | Thr5 | 59.3 | 4.14 | dd | $J_{25,26}$ = 8.6, $J_{26,27}$ = 3.6 |
| 27 | Thr5 | 66.9 | 4.02-3.98 | m | |
| 28 | Thr5 | 19.9 | 1.00 | d | $J_{27,28}$ = 6.3 |
| 29 | Thr5 | 170.0 | -- | -- | |
| 30 | His6 | -- | 7.78 | d | $J_{30,31}$ = 7.5 |
| 31 | His6 | 53.5 | 4.27-4.24 | m | |
| 32 | His6 | 29.2 | 2.97<br>2.82 | dd<br>dd | $J_{32a,32b}$ = 14.8, $J_{31,32a}$ = 5.1<br>$J_{32a,32b}$ = 14.8, $J_{31,32b}$ = 7.4 |
| 33 | His6 | 134.1[a] | -- | -- | |
| 34 | His6 | 117.1[b] | 6.81 | s | |
| 35 | His6 | -- | -- | -- | |
| 36 | His6 | 135.1 | 7.54 | s | |
| 37 | His6 | -- | | -- | |
| 38 | His6 | 173.1 | -- | -- | |

a. $^{13}C$ chemical shifts for His1(4) and His6(33) were obtained from HMBC, and match literature values expected for corresponding carbons in the histidine ring[13]
b. $^{13}C$ chemical shifts for His1(5) and His6(34) were obtained from HMBC and HSQCed spectra, and match literature values expected for corresponding carbons in the histidine ring[13]

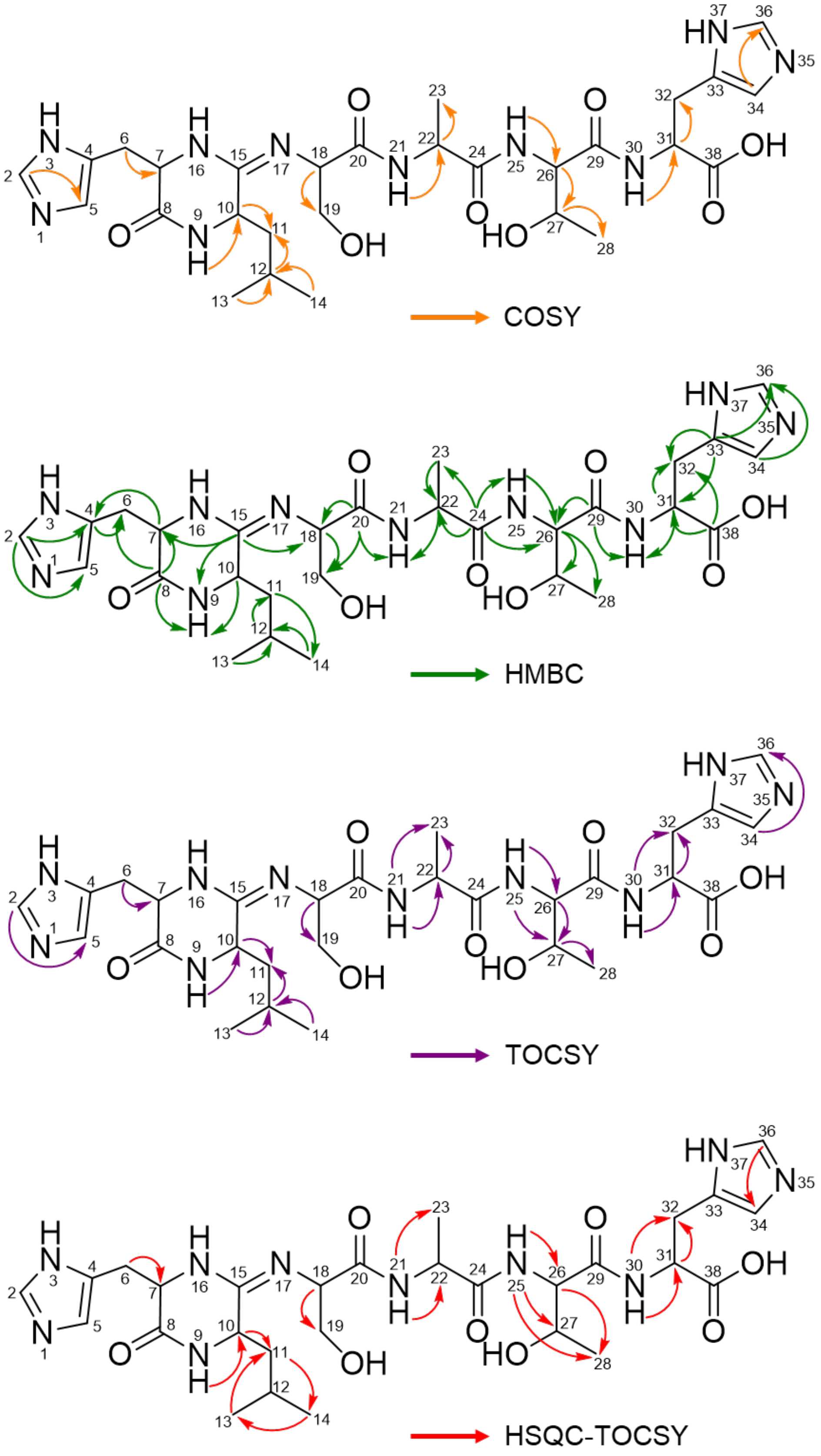

**Figure S6** NMR correlation data observed in 2D spectra (Figures S7 - S14).

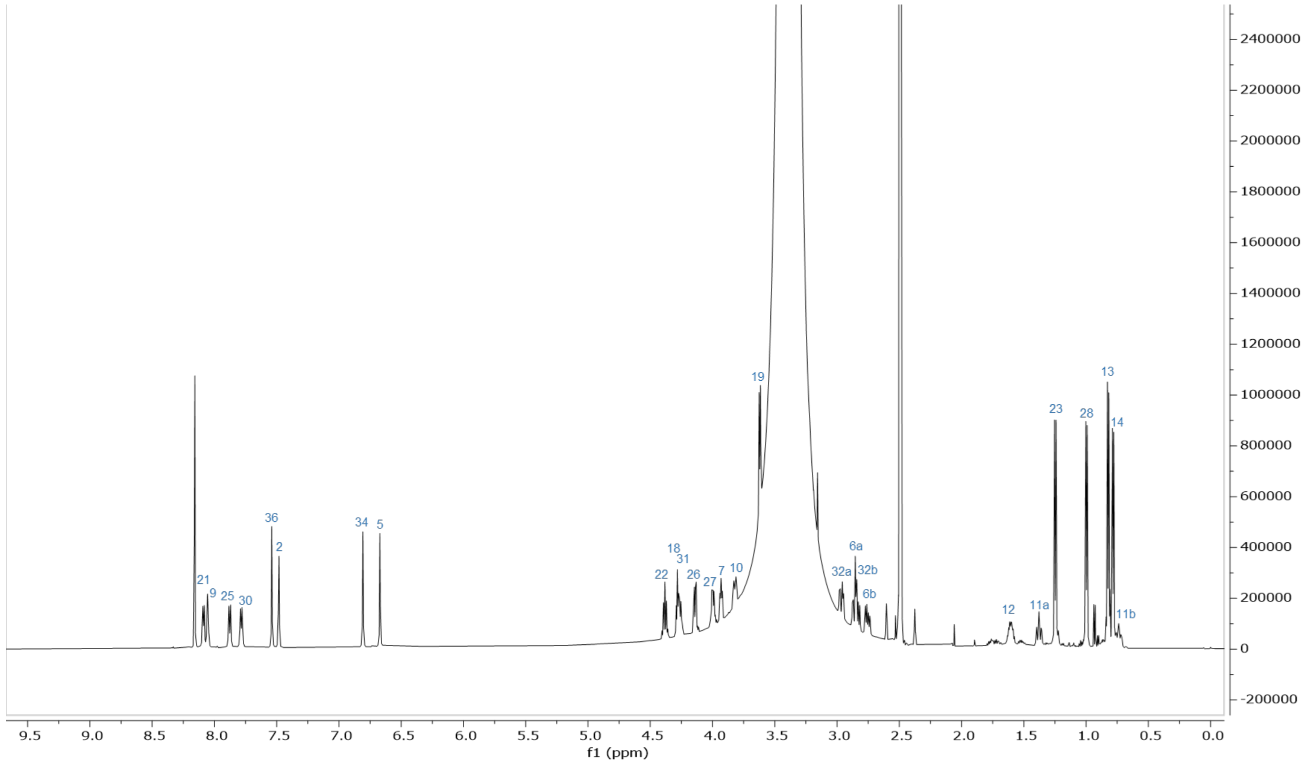

**Figure S7** $^{1}H$ NMR spectrum (600 MHz, DMSO-$d_6$, 298 K) of streptamidine.

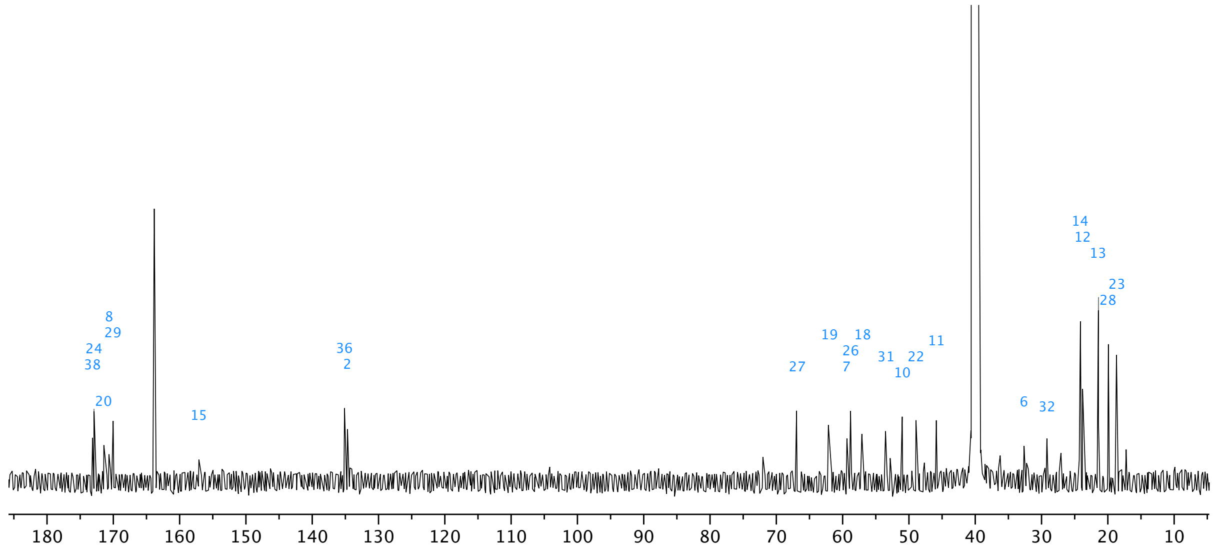

**Figure S8** $^{13}C$ NMR spectrum (150 MHz, DMSO-$d_6$, 298 K) of streptamidine.

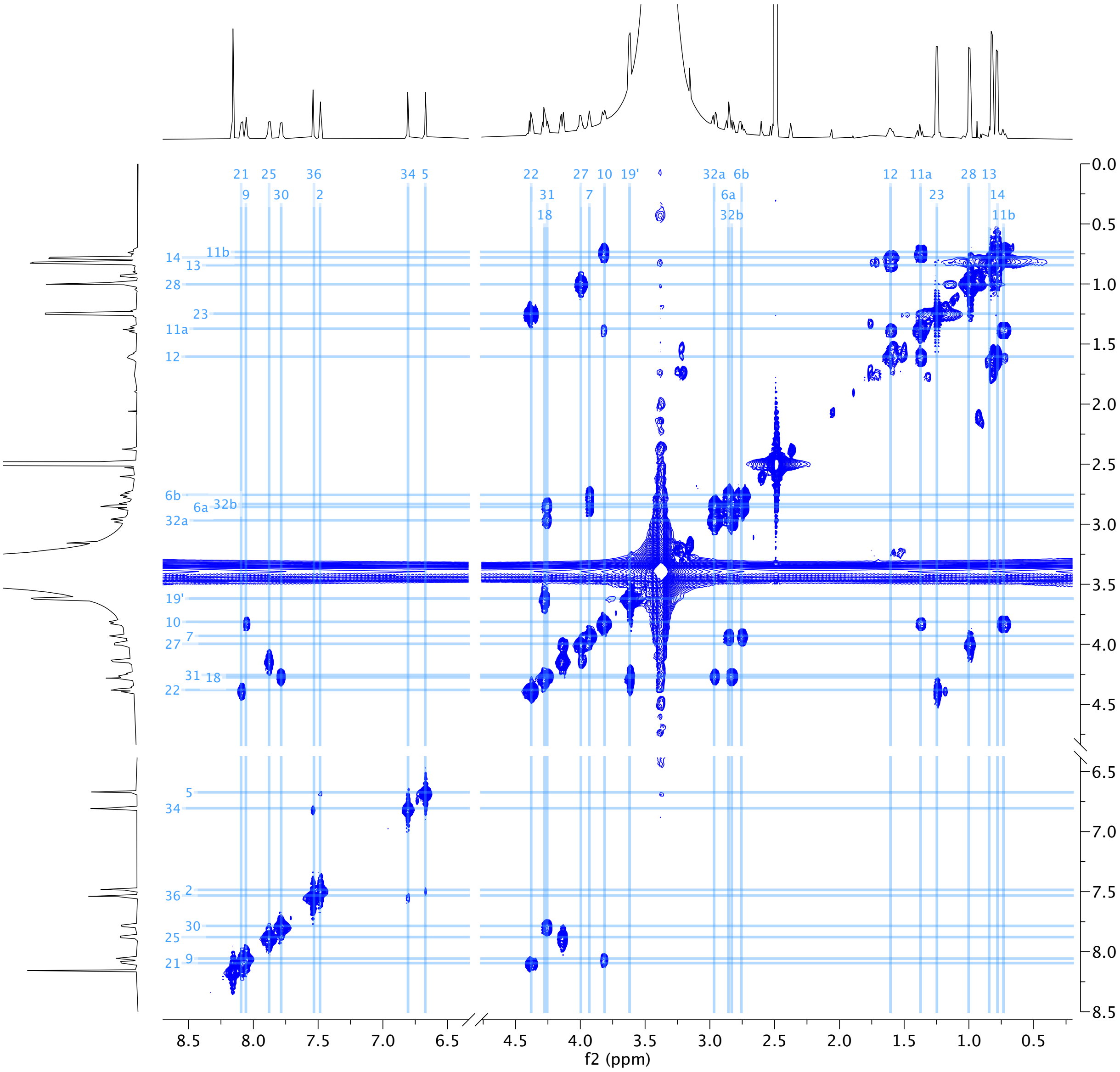

**Figure S9** 2D COSY spectrum (DMSO-$d_6$, 298 K) of streptamidine.

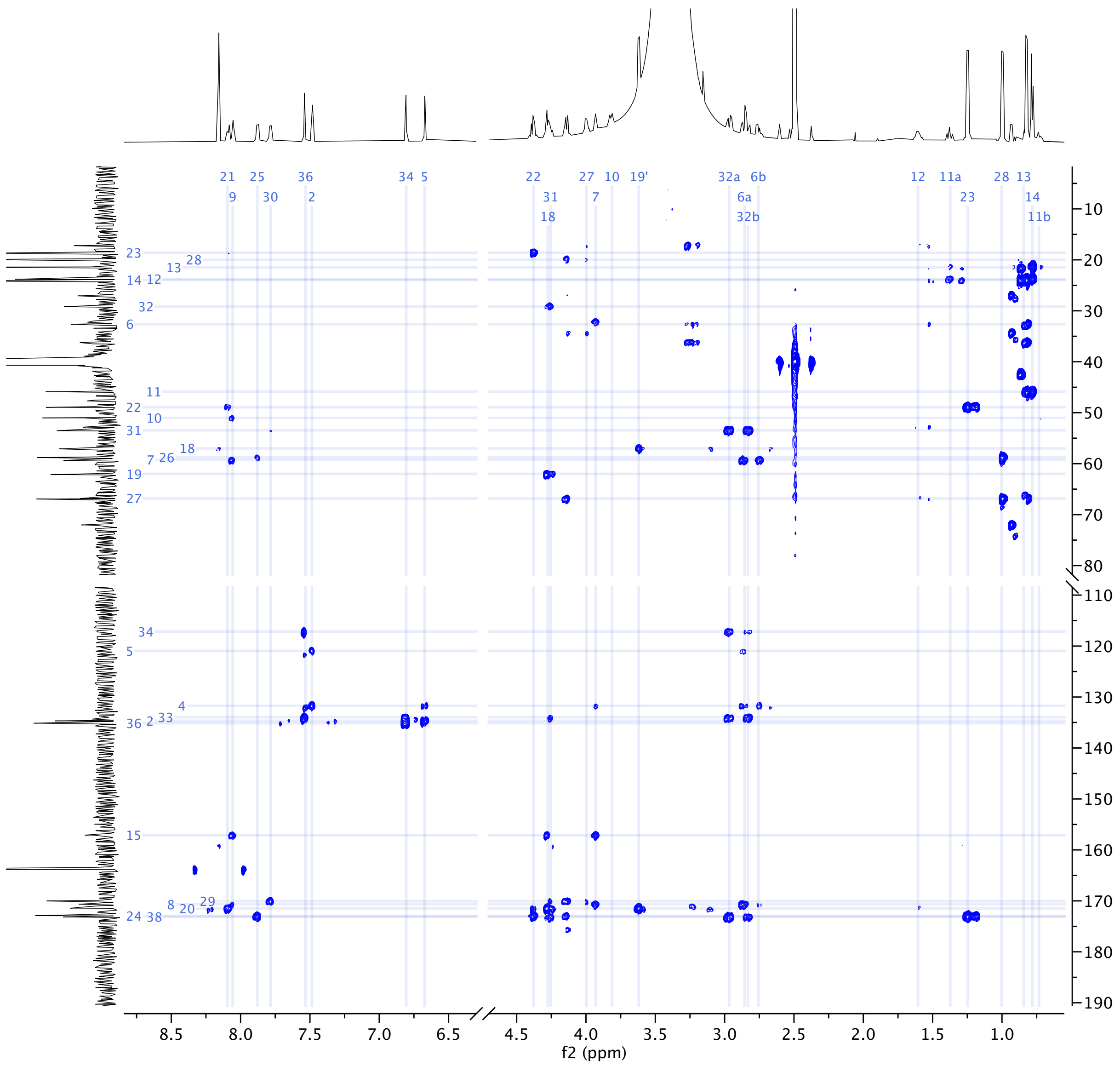

**Figure S10** 2D HMBC spectrum (DMSO-d$_6$, 298 K) of streptamidine.

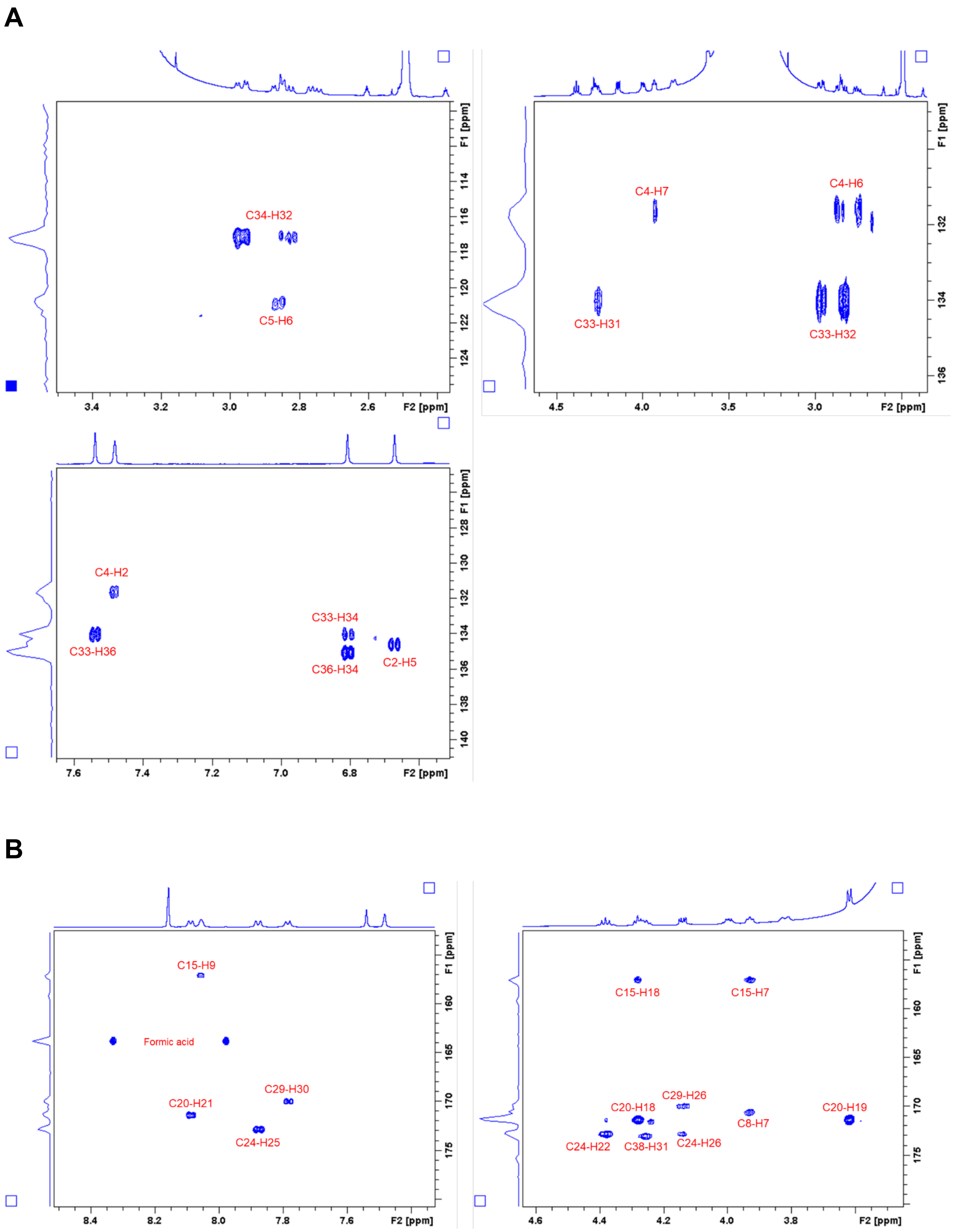

**Figure S11** Selected regions of 2D HMBC spectrum. A. Key correlations for histidine residues. B. Key correlations for amidine carbon and carbonyls.

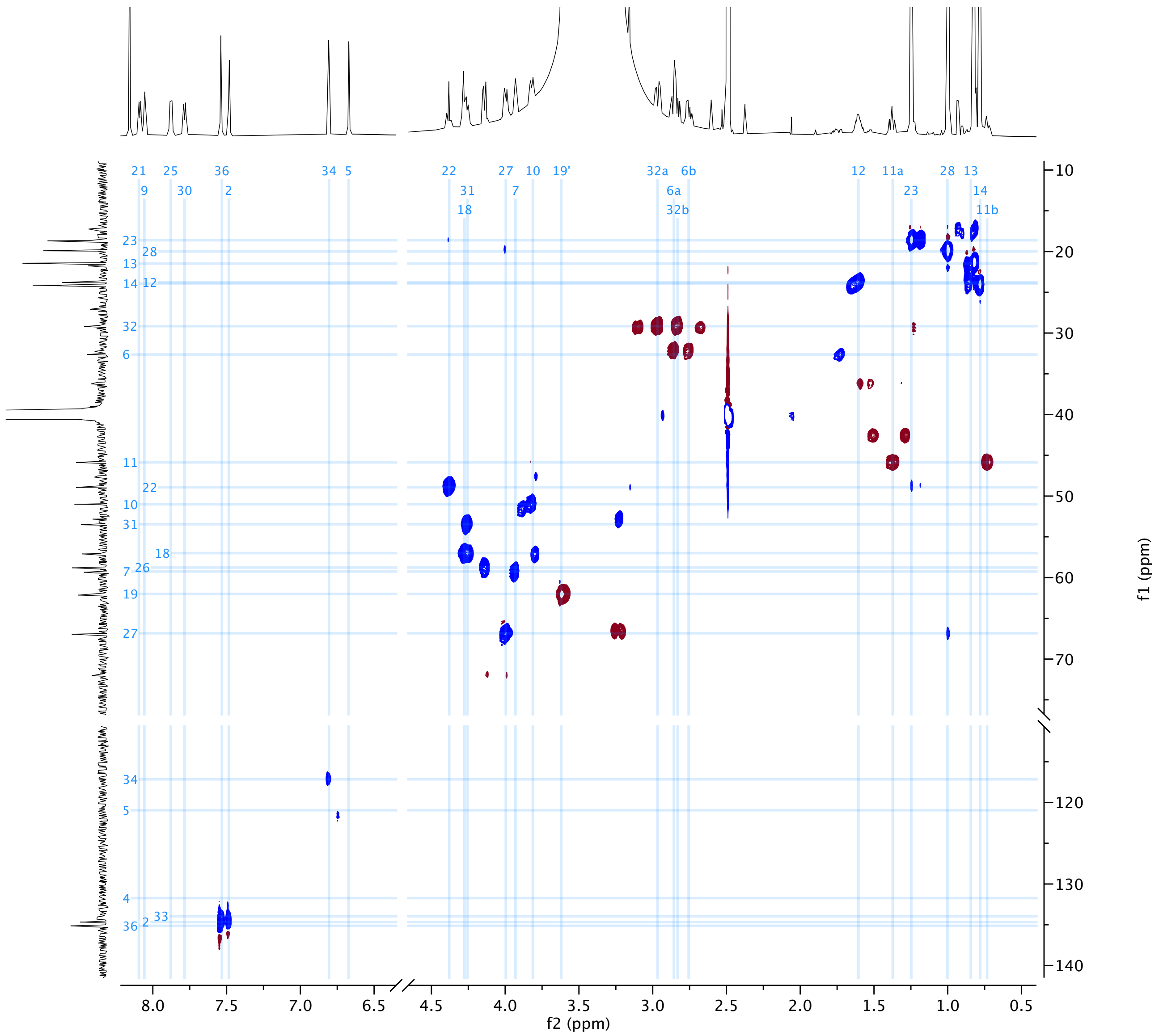

**Figure S12** 2D HSQCed spectrum (DMSO-$d_6$, 298 K) of streptamidine.

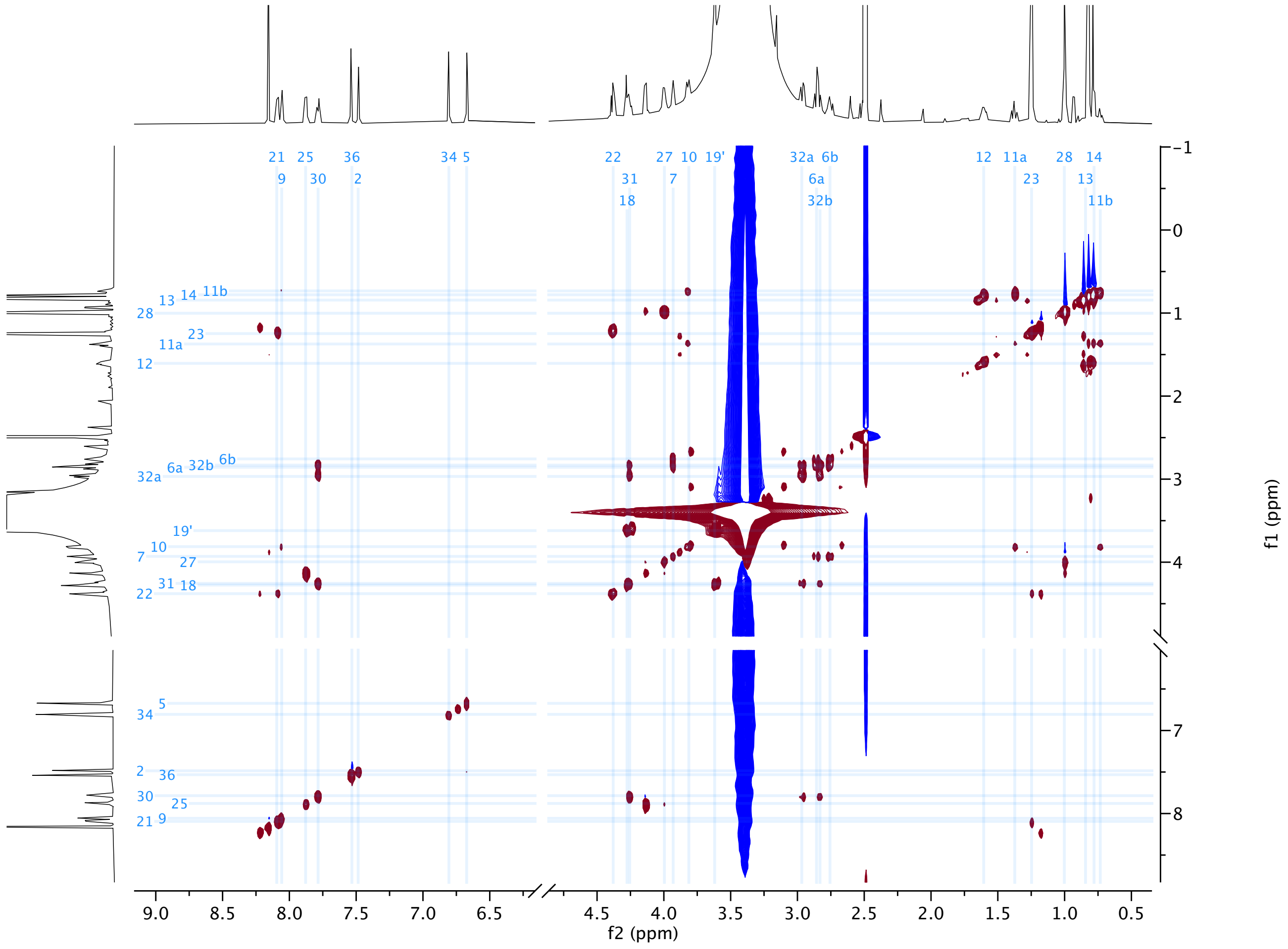

**Figure S13** 2D TOCSY spectrum (DMSO-d$_6$, 298 K) of streptamidine.

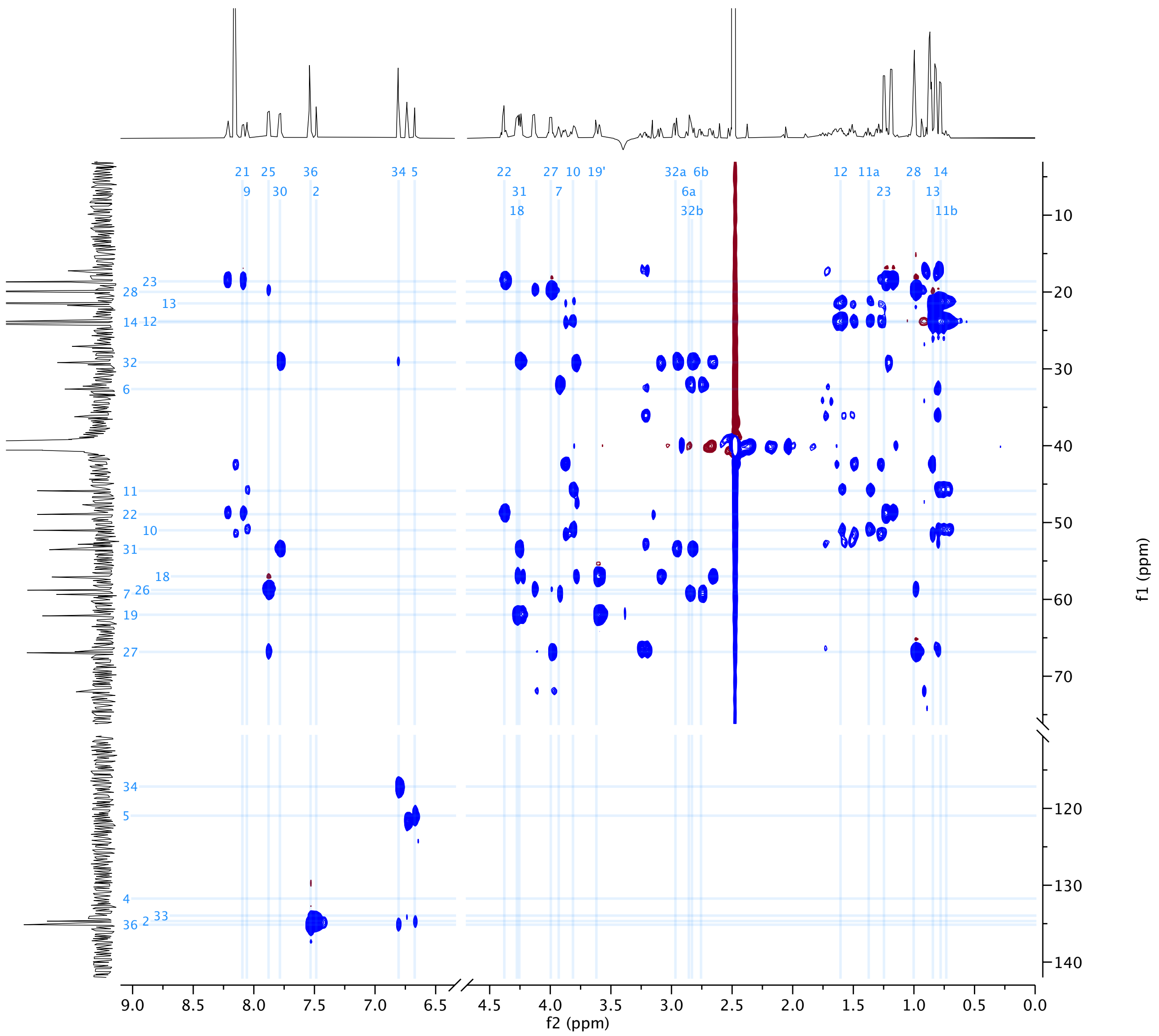

**Figure S14** 2D HSQC-TOCSY spectrum (DMSO-d$_6$, 298 K) of streptamidine. Some extra cross-peaks (e.g. for C5) are predicted to relate to an isomerisation during the acquisition of spectra.

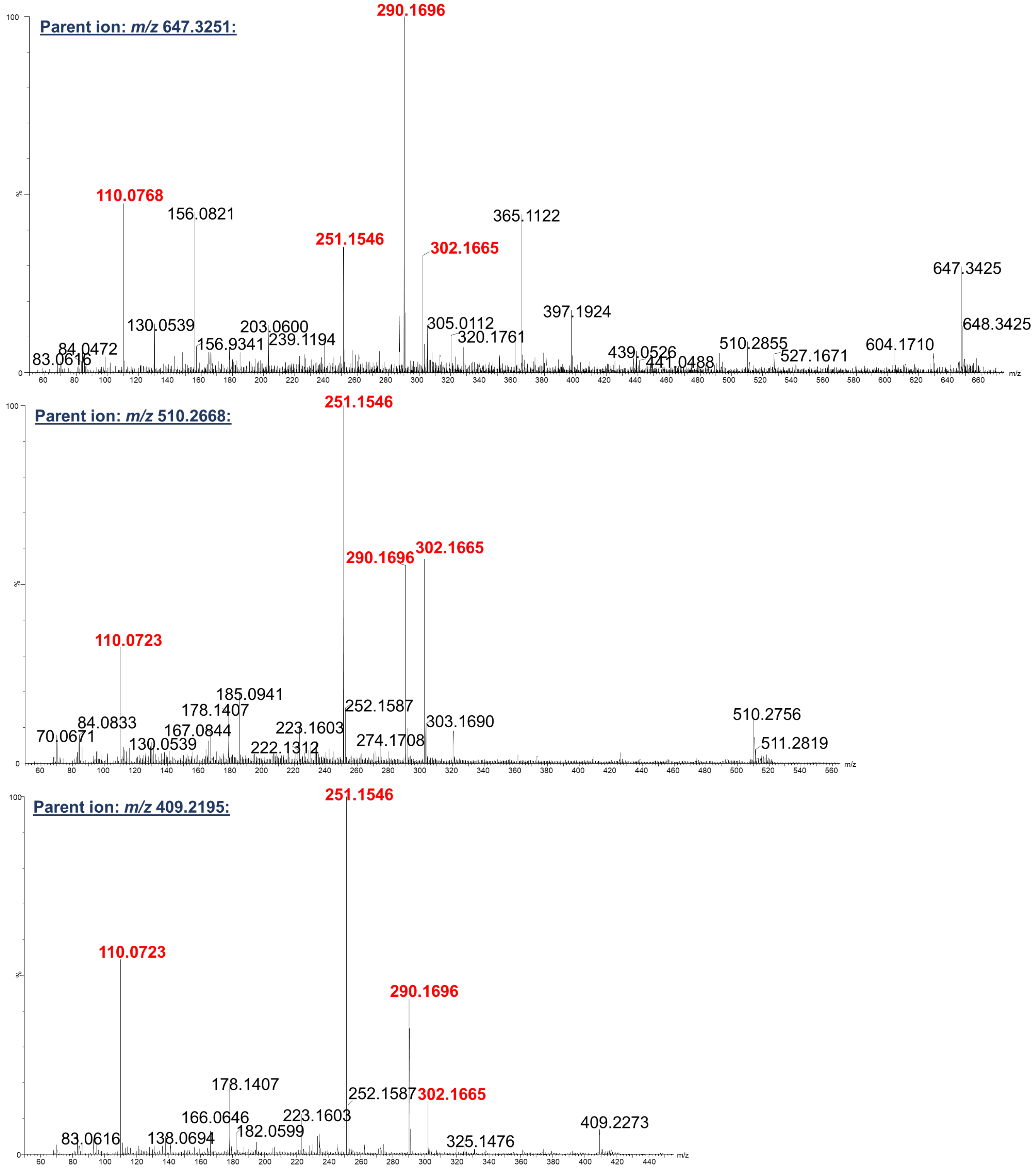

**Figure S15** MS$^2$ fragmentation data for streptamidine (647.3251), predicted modified HLSAT (510.2668) and predicted modified HLSA (409.2195), obtained using a Waters Synapt G2Si.

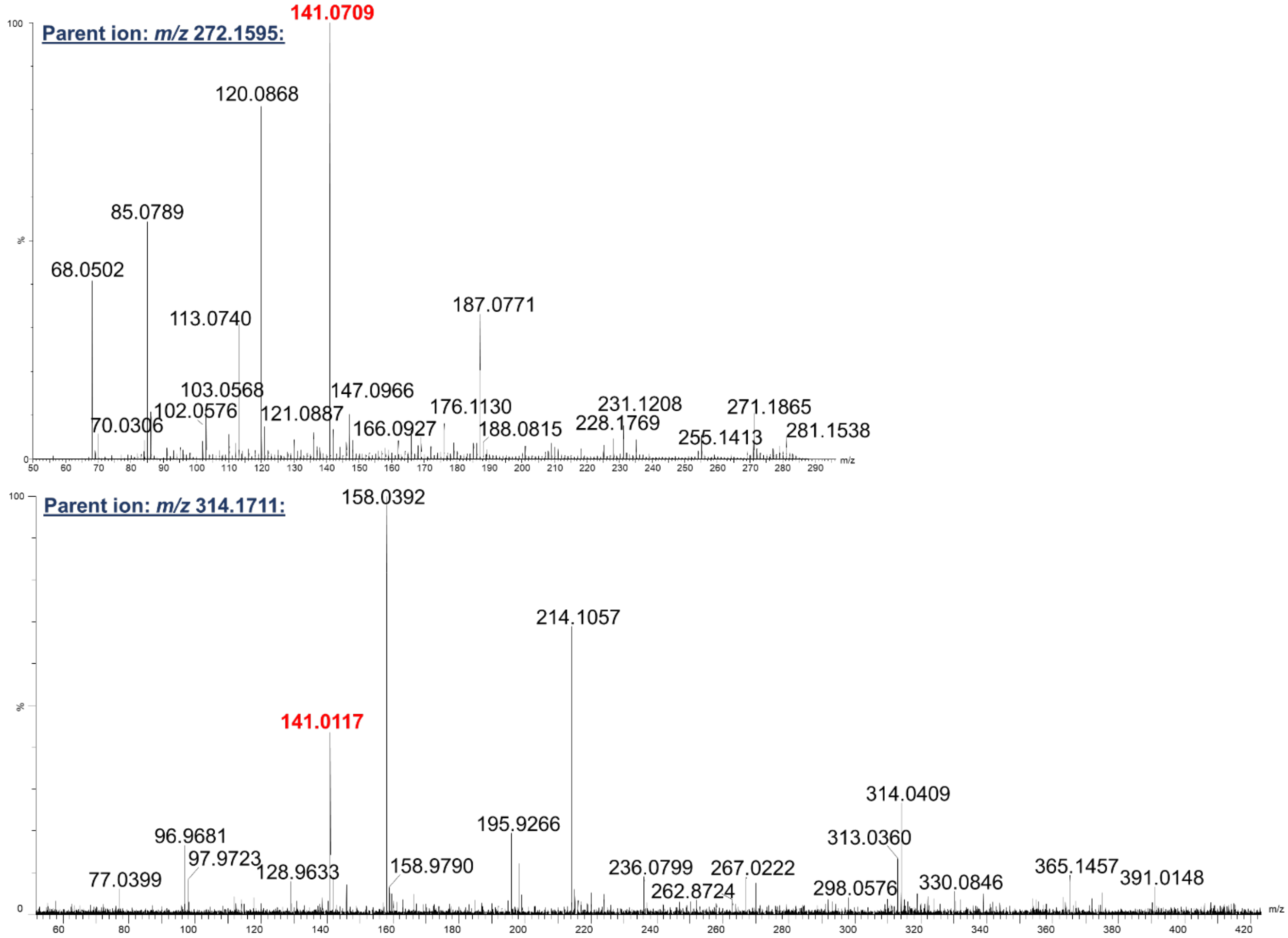

**Figure S16** MS$^2$ fragmentation data for predicted dehydrated LSA (272.1595) and predicted acetylated and dehydrated LSA (314.1711), obtained using a Waters Synapt G2Si.

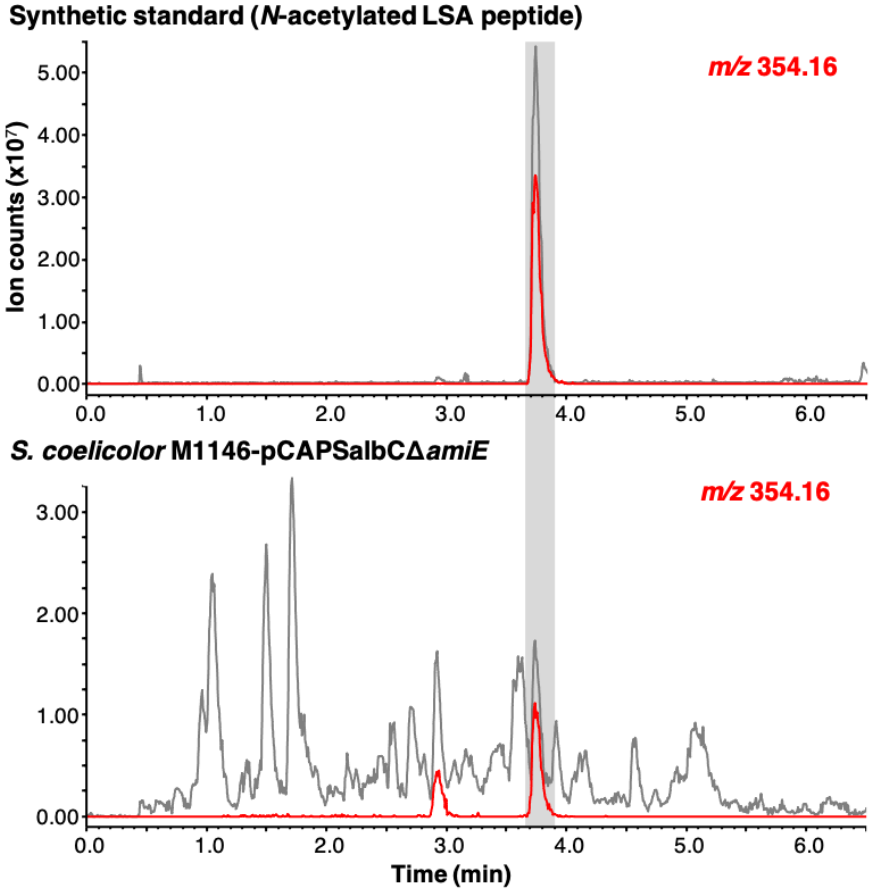

**Figure S17** LC-MS analysis (Shimadzu IT-TOF) of *S. coelicolor* M1146-SalbC Δ*amiE* culture extract versus a synthetic standard of *N*-acetylated LSA peptide (grey line = base peak chromatogram). Extracted ion chromatogram of *m/z* 354.16 (red) indicates expected mass of sodium adduct of *N*-acetylated LSA. This mass elutes in both samples at ~3.75 min, indicating that this same peptide is produced by the dehydrogenase mutant.

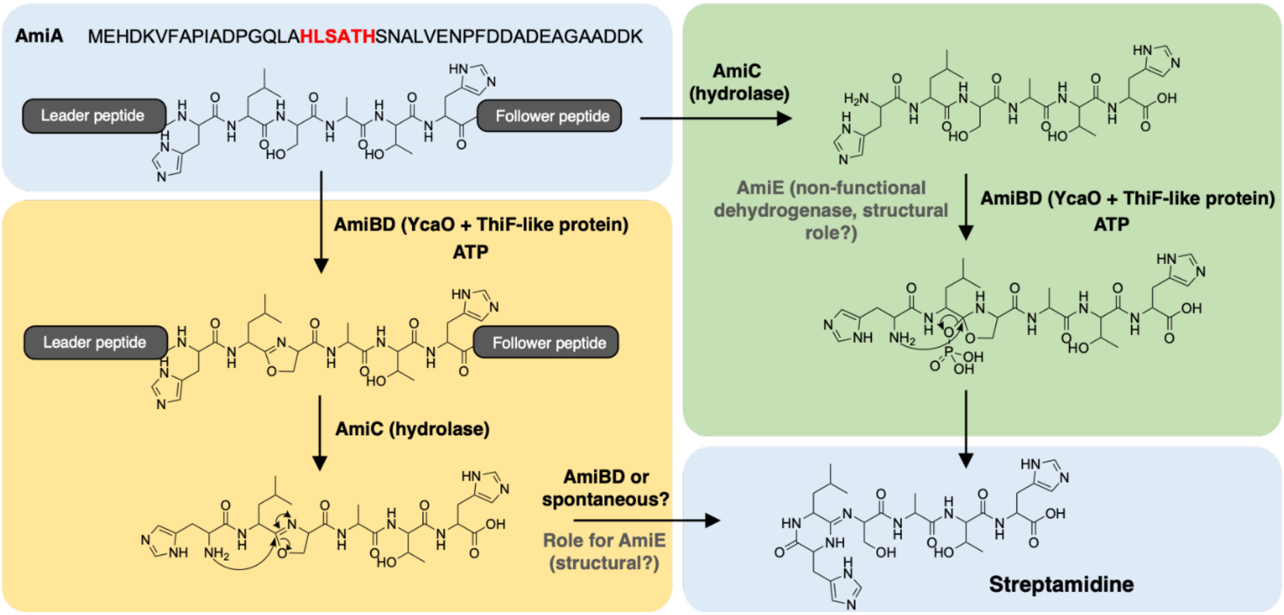

**Figure S18** Proposed mechanism of amidine ring formation in streptamidine based on predicted intermediates produced by dehydrogenase mutant. The yellow box represents a route that goes via a stable oxazoline intermediate prior to leader/follower peptide removal, where AmiE potentially has a structural role within an AmiBDE complex for proper amidine formation but is not needed for initial ATP-dependent oxazoline formation. The green box represents a route where the leader/follower peptide is removed first, thereby providing a free *N*-terminal amine for cyclisation via an *O*-phosphorylated hemiorthoamide. In the absence of AmiE, the final amidine forming step could potentially be disrupted.

**Table S10** Metabolic data of complete TAR clone (SalbC) expressed in *S. coelicolor* M1146 versus precursor peptide mutant (SalbCΔP) and medium only (SM12). Analysis carried out using Profiling Solution (Shimadzu) where numbers reflect peak areas for specific metabolites.

| Ion m/z | Ion RT | SalbC | SalbC | SalbC | SalbC | SalbCΔP | SalbCΔP | SalbCΔP | SalbCΔP | SM12 |
|---|---|---|---|---|---|---|---|---|---|---|
| 510.2674 | 1.09 | 2499654 | 2564489 | 1922639 | 2372999 | 0 | 0 | 0 | 0 | 0 |
| 324.1662 | 0.877 | 2207472 | 2081116 | 2825998 | 2730479 | 0 | 0 | 0 | 0 | 0 |
| 409.2172 | 1.012 | 1595479 | 1643531 | 2001746 | 1911645 | 0 | 0 | 0 | 0 | 204194 |
| 272.1603 | 2.423 | 996032 | 865128 | 846665 | 990778 | 0 | 0 | 0 | 0 | 0 |
| 314.1695 | 2.48 | 873422 | 733426 | 962465 | 1160814 | 0 | 0 | 0 | 0 | 0 |
| 338.1806 | 0.782 | 864190 | 699589 | 767195 | 924738 | 0 | 0 | 0 | 0 | 0 |
| 647.324 | 0.869 | 546123 | 492537 | 588600 | 703503 | 0 | 0 | 0 | 0 | 0 |
| 332.1807 | 2.475 | 385982 | 344655 | 416676 | 487723 | 0 | 0 | 0 | 0 | 0 |
| 619.5266 | 7.746 | 8982193 | 9390319 | 9782771 | 9933121 | 9173293 | 9540028 | 9587654 | 9905776 | 8688859 |
| 381.2967 | 6.984 | 8167554 | 8251210 | 7813885 | 8525158 | 7619021 | 8403720 | 8255679 | 7662858 | 6411262 |
| 647.5576 | 7.102 | 6323495 | 6894933 | 7087683 | 7531915 | 6990360 | 7383752 | 7449997 | 7319537 | 7779696 |
| 591.4958 | 7.476 | 4950153 | 5169159 | 5696559 | 5770050 | 5196230 | 5368502 | 5705355 | 5799733 | 4599594 |
| 365.1068 | 0.388 | 4935655 | 4497150 | 4589359 | 4819098 | 5041421 | 4753954 | 4769492 | 5080979 | 540055 |
| 561.3606 | 3.235 | 4637774 | 5156793 | 5881030 | 6046490 | 4854390 | 5864654 | 5646551 | 5734262 | 0 |
| 623.3361 | 4.181 | 4591141 | 5027047 | 5267581 | 5132072 | 4250342 | 4635180 | 4487253 | 4448017 | 0 |
| 614.2726 | 2.347 | 3941212 | 2528841 | 2196621 | 1601265 | 3066789 | 2042013 | 1532947 | 1411457 | 0 |
| 353.2662 | 6.933 | 3931686 | 4030330 | 3894694 | 4267148 | 3810405 | 4044643 | 3995562 | 3885436 | 3047360 |
| 257.1473 | 0.371 | 3680536 | 3594810 | 3954374 | 3806560 | 3582185 | 3837819 | 3836813 | 3302630 | 2457808 |

**Table S11** Metabolic data of *S. albus* J1074 wild type versus pathway-disrupted mutant and medium only (SM12). Analysis carried out as described for Table S10.

| Ion m/z | Ion RT | *S. albus* | *S. albus* | *S. albus* | *S. albus* | Mutant | Mutant | Mutant | Mutant | SM12 |
|---|---|---|---|---|---|---|---|---|---|---|
| 272.15 | 3.2 | 0 | 0 | 0 | 0 | 157702 | 217032 | 239481 | 157170 | 0 |
| 320.166 | 1.260 | 197786 | 251664 | 260141 | 225336 | 0 | 0 | 0 | 0 | 0 |
| 338.177 | 1.352 | 959988 | 1359900 | 989904 | 994114 | 0 | 108668 | 140099 | 131368 | 0 |
| 510.260 | 3.152 | 111700 | 646627 | 552968 | 108028 | 95654 | 0 | 0 | 0 | 0 |
| 510.266 | 2.521 | 267676 | 309829 | 307114 | 259442 | 0 | 0 | 0 | 0 | 0 |
| 647.327 | 1.738 | 372300 | 544327 | 573868 | 484952 | 0 | 0 | 0 | 0 | 0 |
| 200.0412 | 0.449 | 202261 | 171648 | 178763 | 187584 | 148285 | 138368 | 186912 | 124928 | 0 |
| 200.0855 | 4.958 | 0 | 0 | 0 | 0 | 0 | 0 | 98816 | 0 | 0 |
| 200.0879 | 3.431 | 123351 | 0 | 0 | 0 | 93245 | 0 | 171980 | 101888 | 0 |
| 200.0885 | 3.642 | 0 | 0 | 0 | 0 | 0 | 156142 | 0 | 0 | 0 |
| 201.1149 | 3.4 | 0 | 0 | 0 | 0 | 0 | 0 | 94744 | 145183 | 0 |
| 201.1165 | 3.529 | 102016 | 0 | 85312 | 133288 | 148892 | 0 | 107375 | 146578 | 0 |
| 201.1175 | 4.948 | 115264 | 0 | 0 | 104000 | 85376 | 0 | 110193 | 0 | 0 |
| 201.1184 | 5.823 | 0 | 0 | 0 | 105417 | 222258 | 102144 | 0 | 78912 | 0 |
| 201.119 | 3.671 | 0 | 0 | 0 | 0 | 192297 | 109793 | 0 | 0 | 0 |
| 201.1197 | 3.777 | 0 | 141010 | 0 | 211287 | 0 | 138496 | 128204 | 161206 | 0 |
| 201.1199 | 5.479 | 932083 | 988136 | 1020146 | 1214868 | 1009633 | 1008262 | 984542 | 1033656 | 0 |

**Table S12** Metabolic data of complete TAR clone (SalbC) expressed in *S. coelicolor* M1146 versus pathway mutants (Δ*amiB*, Δ*amiC*, Δ*amiD*, Δ*amiE*, Δ*amiX*) and medium only (SM12). Analysis carried out as described for Table S10.

| Ion m/z | Ion RT | SalbC | SalbC | ΔamiB | ΔamiB | ΔamiB | ΔamiC | ΔamiC | ΔamiC | ΔamiD | ΔamiD | ΔamiD | ΔamiE | ΔamiE | ΔamiE | ΔamiX | ΔamiX | ΔamiX | SM12 1 | SM12 2 |
|---|---|---|---|---|---|---|---|---|---|---|---|---|---|---|---|---|---|---|---|---|
| 324.1659 | 1.398 | 2262279 | 2214873 | 0 | 0 | 0 | 0 | 0 | 0 | 0 | 0 | 0 | 0 | 0 | 0 | 1057241 | 1172300 | 1139022 | 0 | 0 |
| 510.2698 | 1.55 | 1475999 | 1171449 | 0 | 0 | 0 | 0 | 0 | 0 | 0 | 0 | 0 | 0 | 0 | 0 | 810404 | 763166 | 660021 | 0 | 0 |
| 314.1696 | 3.025 | 1042037 | 847530 | 0 | 0 | 0 | 0 | 0 | 0 | 0 | 0 | 0 | 1605224 | 2048790 | 1588020 | 2825272 | 2569195 | 2451654 | 0 | 0 |
| 647.3248 | 1.392 | 544662 | 545338 | 0 | 0 | 0 | 0 | 0 | 0 | 0 | 0 | 0 | 0 | 0 | 0 | 290000 | 237703 | 223171 | 0 | 0 |
| 332.1785 | 3.023 | 393459 | 329963 | 0 | 0 | 0 | 0 | 0 | 0 | 0 | 0 | 0 | 887866 | 1116939 | 712023 | 1980880 | 1813192 | 1386092 | 0 | 0 |
| 272.1582 | 2.983 | 387710 | 410005 | 0 | 0 | 0 | 0 | 0 | 0 | 0 | 0 | 0 | 796578 | 1351619 | 943842 | 1570774 | 1699116 | 1560029 | 0 | 0 |
| 338.1794 | 0.96 | 349530 | 367394 | 0 | 0 | 0 | 0 | 0 | 0 | 0 | 0 | 0 | 0 | 0 | 0 | 200116 | 191623 | 239485 | 141566 | 131291 |
| 354.1602 | 3.034 | 280038 | 252484 | 0 | 0 | 0 | 0 | 0 | 0 | 0 | 0 | 0 | 443885 | 483075 | 389179 | 649966 | 604358 | 585720 | 0 | 0 |
| 510.2688 | 1.202 | 205147 | 197976 | 0 | 0 | 0 | 0 | 0 | 0 | 0 | 0 | 0 | 0 | 0 | 0 | 172142 | 149342 | 169857 | 0 | 0 |

**Table S13** Metabolic data of complete TAR clone (SalbC) expressed in *S. coelicolor* M1146 versus pathway mutants: iron transporter deletion (IrTr), oxygenase deletion (Oxy), peptide methionine sulfoxide reductase deletion (PM) MarR deletion (MarR), ABC transporter deletion (ABC), acetyltransferase deletion (AT) and medium only (SM12). Analysis carried out as described for Table S10.

| Ion m/z | Ion RT | SalbC | SalbC | SalbC | IrTr | IrTr | IrTr | Oxy | Oxy | Oxy | PM | PM | PM | MarR | MarR | MarR | ABC | ABC | ABC | AT | AT | AT |
|---|---|---|---|---|---|---|---|---|---|---|---|---|---|---|---|---|---|---|---|---|---|---|
| 324.1652 | 1.335 | 2262502 | 2715880 | 2252375 | 0 | 0 | 0 | 2638215 | 3555376 | 141632 | 3046506 | 2325230 | 2642271 | 3931198 | 2664038 | 4194032 | 389484 | 206483 | 0 | 2608474 | 129689 | 0 |
| 200.0433 | 0.737 | 0 | 0 | 0 | 403997 | 99072 | 0 | 107414 | 0 | 283233 | 0 | 72576 | 0 | 0 | 0 | 0 | 238197 | 296059 | 234648 | 224119 | 234583 | 0 |
| 201.1138 | 2.69 | 0 | 0 | 0 | 0 | 0 | 0 | 0 | 0 | 0 | 0 | 0 | 0 | 0 | 0 | 0 | 0 | 0 | 0 | 125322 | 0 | 0 |
| 201.1184 | 3.907 | 0 | 0 | 0 | 0 | 0 | 0 | 0 | 0 | 0 | 0 | 0 | 0 | 0 | 0 | 0 | 0 | 0 | 0 | 142073 | 0 | 0 |
| 209.1259 | 1.328 | 0 | 0 | 0 | 63639 | 0 | 88353 | 0 | 0 | 0 | 0 | 0 | 0 | 0 | 0 | 0 | 0 | 0 | 69120 | 0 | 110878 | 0 |
| 209.1316 | 1.255 | 0 | 0 | 0 | 0 | 0 | 0 | 0 | 0 | 0 | 0 | 0 | 0 | 0 | 0 | 0 | 0 | 0 | 0 | 0 | 0 | 78303 |
| 210.1147 | 1.227 | 0 | 0 | 0 | 0 | 0 | 59392 | 0 | 0 | 0 | 0 | 0 | 68141 | 0 | 89269 | 0 | 0 | 0 | 0 | 0 | 0 | 0 |
| 210.1171 | 1.31 | 0 | 0 | 0 | 0 | 0 | 0 | 0 | 0 | 0 | 0 | 0 | 0 | 0 | 0 | 0 | 0 | 0 | 73833 | 0 | 0 | 0 |
| 212.107 | 1.293 | 0 | 0 | 0 | 105723 | 116932 | 128830 | 0 | 0 | 89186 | 0 | 0 | 0 | 0 | 0 | 0 | 0 | 119488 | 112772 | 0 | 92070 | 77554 |
| 213.1209 | 3.19 | 95168 | 119609 | 96907 | 0 | 72384 | 0 | 110988 | 92032 | 69312 | 65920 | 85622 | 68330 | 82432 | 88704 | 94740 | 98816 | 78976 | 56128 | 101287 | 59328 | 0 |
| 213.1224 | 3.07 | 0 | 0 | 0 | 188868 | 198530 | 80000 | 0 | 0 | 0 | 0 | 0 | 78528 | 0 | 0 | 0 | 0 | 0 | 0 | 0 | 0 | 0 |
| 215.1365 | 3.294 | 294120 | 158151 | 258419 | 417093 | 470743 | 493433 | 0 | 0 | 308759 | 175334 | 273194 | 310409 | 0 | 201804 | 0 | 128783 | 187794 | 412791 | 0 | 357362 | 342869 |
| 215.1381 | 4.111 | 0 | 0 | 0 | 0 | 0 | 0 | 87419 | 0 | 0 | 72640 | 96585 | 75840 | 75814 | 66368 | 0 | 0 | 88896 | 0 | 0 | 0 | 0 |

**Table S14** Indicator strains tested in bioactivity assays.

| **Organism tested** | **Description** | **Inhibition observed** |
|---|---|---|
| *Escherichia coli* ATCC25922 | Gram-negative bacterium | None |
| *E. coli* NR986 | Gram-negative bacterium (mutant with increased membrane permeability)[14] | None |
| *Pseudomonas aeruginosa* PA01 | Gram-negative bacterium | None |
| *Pseudomonas fluorescens* | Gram-negative bacterium | None |
| *Bacillus subtilis* 168 | Gram-positive bacterium | None |
| *Micrococcus luteus* | Gram-positive bacterium | None |
| *Mycobacterium smegmatis* MC2155 | Gram-positive bacterium | None |
| *Streptomyces scabies* | Gram-positive bacterium | None |
| *Streptomyces cattleya* | Gram-positive bacterium | None |
| *Candida utilis* | Fungus | None |

**Figure S19 (shown below)** Alignment of precursor peptides from network 1 containing motif A. Aligned with MEGA7 software.[15]

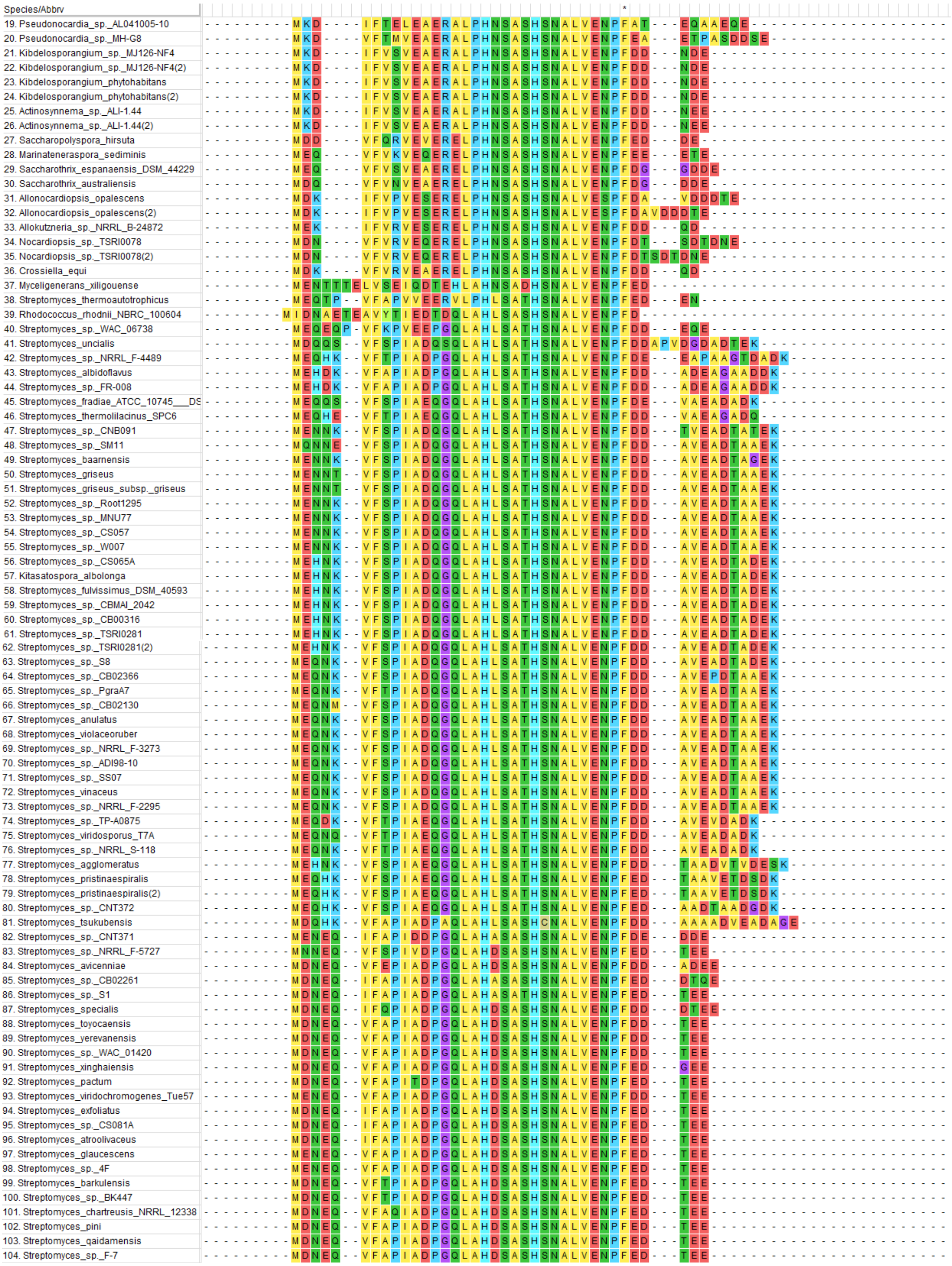

### Figure S19 continued

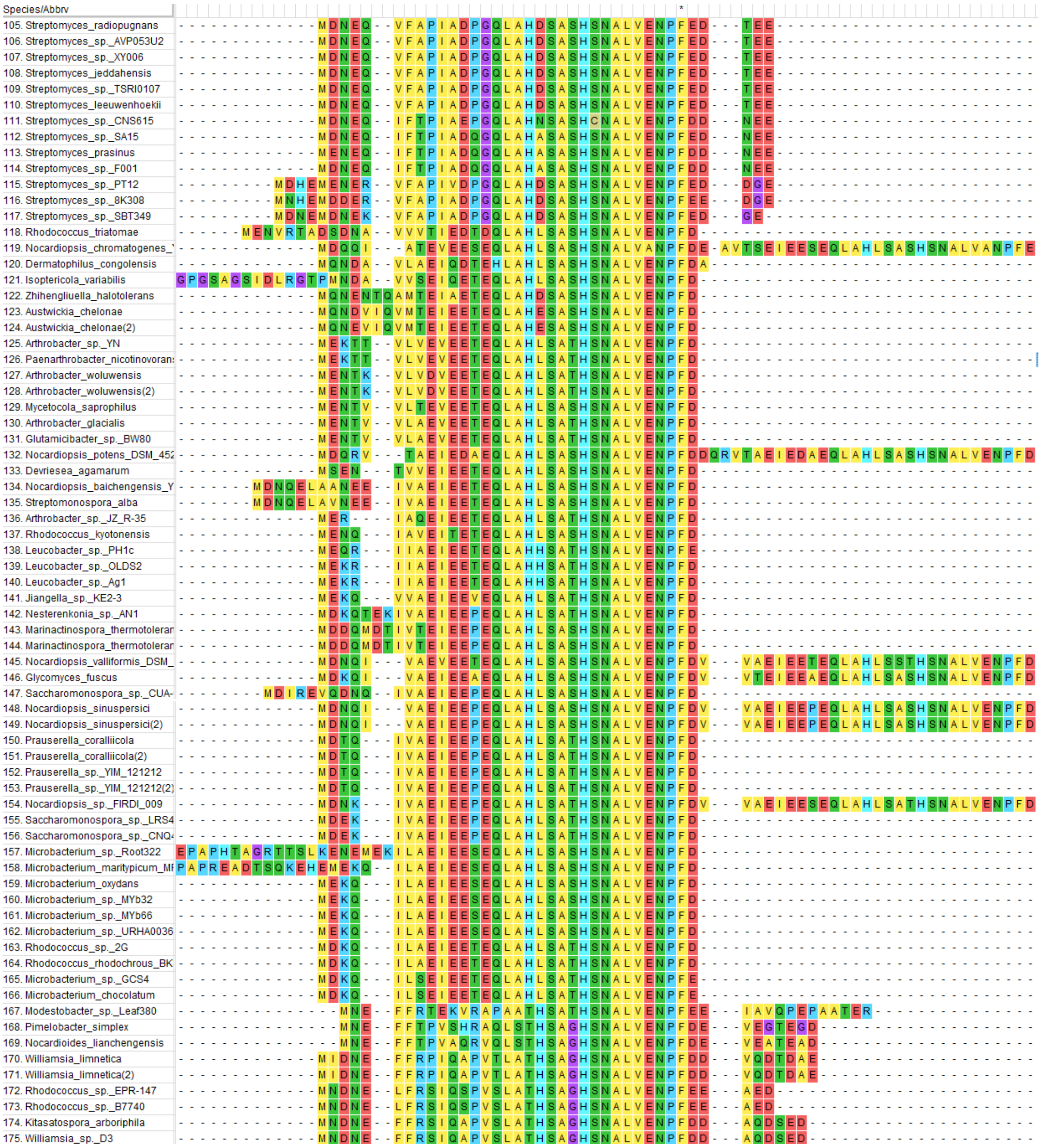

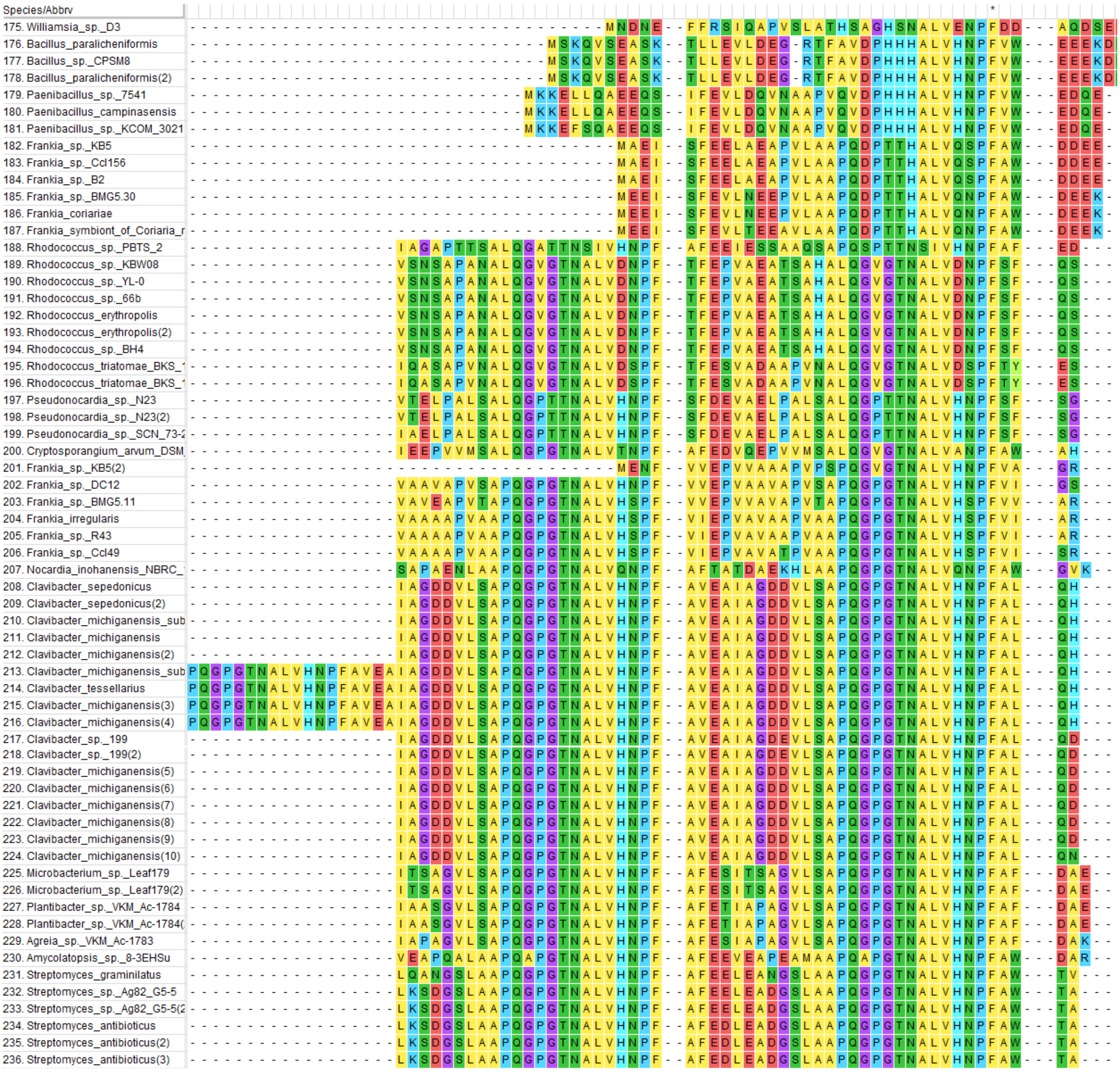

**Figure S20** Alignment of precursor peptides from network 1 containing motif B. Aligned with MEGA7 software.

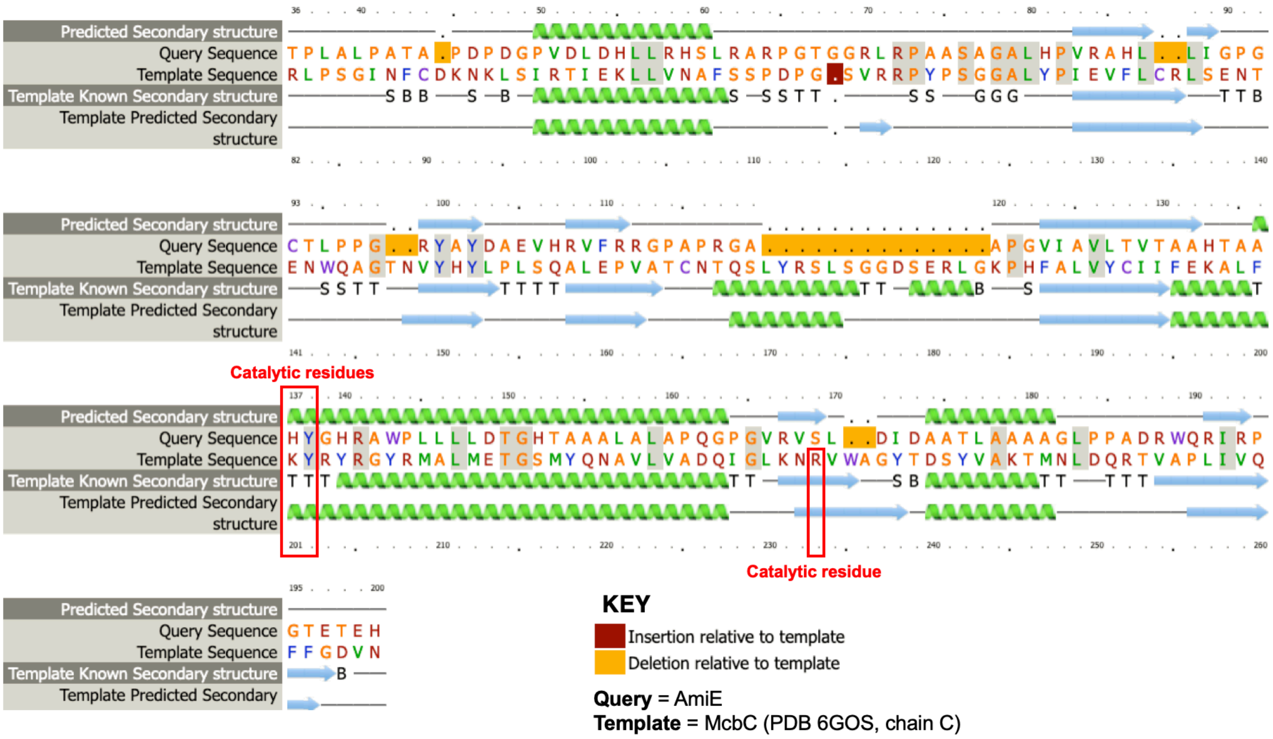

**Figure S21** Secondary structure alignment of AmiE with McbC (PDB 6GOS) from the microcin B17 pathway[16] generated with Phyre2.[17] Catalytic residues identified by Ghilarov *et al.*[16] in McbC are highlighted.

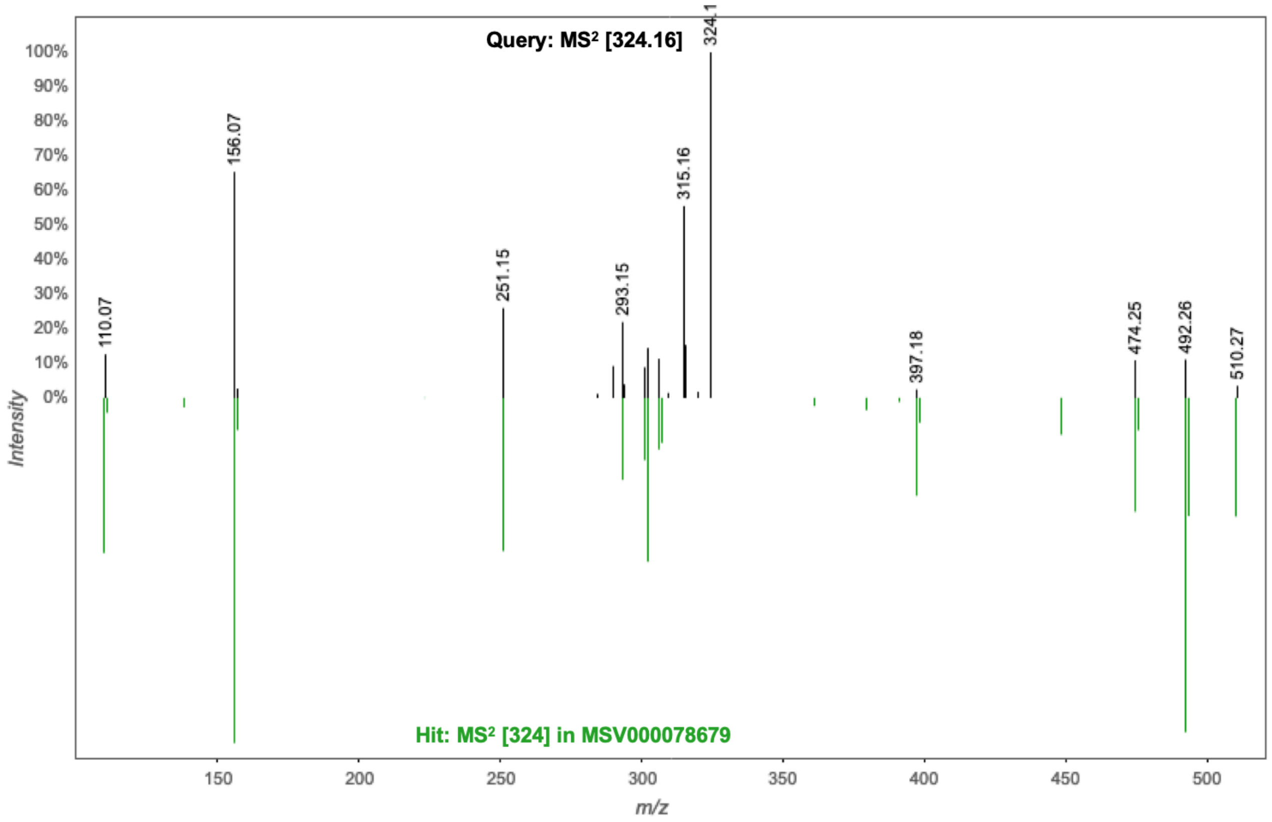

**Figure S22** Spectral match to the MS$^2$ spectrum of [streptamidine+2H]$^{2+}$ (*m/z* 324.16) identified using MASST (Mass Spectrometry Search Tool)[18] at GNPS (Global Natural Products Social Molecular Networking). The hit is found in multiple samples of MassIVE Dataset MSV000078679 (“Zhang lab_microbes library_MS130001~9”), which is defined as an actinomycete dataset. The non-matching 324.1 relates to the unfragmented parent ion in the query spectrum.
